# A genetically adaptable strategy for ribose scavenging in a human gut symbiont plays a diet-dependent role in colon colonization

**DOI:** 10.1101/574913

**Authors:** Robert W. P. Glowacki, Nicholas A. Pudlo, Yunus Tuncil, Ana S. Luis, Anton I. Terekhov, Bruce R. Hamaker, Eric C. Martens

**Affiliations:** Department of Microbiology and Immunology, University of Michigan Medical School, Ann Arbor, MI 48109; Department of Food Science and Whistler Center for Carbohydrate Research, Purdue University, West Lafayette, IN 47907; Department of Food Engineering, Ordu University, Ordu, Turkey

## Abstract

Efficient nutrient acquisition in the competitive human gut is essential for microbial persistence. While polysaccharides have been well-studied nutrients for the gut microbiome, other resources such as co-factors and nucleic acids have been less examined. We describe a series of ribose utilization systems (RUSs) that are broadly represented in Bacteroidetes and appear to have diversified to allow access to ribose from a variety of substrates. One *Bacteroides thetaiotaomicron* RUS variant is critical for competitive gut colonization in a diet-specific fashion. Using molecular genetics, we probed the nature of the ribose source underlying this diet-specific phenotype, revealing that hydrolytic functions in RUS (*e.g*., to cleave ribonucleosides) are present but dispensable. Instead, ribokinases that are activated *in vivo* and participate in cellular ribose-phosphate metabolism are essential. Our results underscore the extensive mechanisms that gut symbionts have evolved to access nutrients and how metabolic context determines the impact of these functions *in vivo*.

## Introduction

Symbiotic microorganisms that inhabit the human intestine complement digestive capacity in numerous ways, with the most mechanistically understood examples involving degradation and fermentation of chemically diverse fiber polysaccharides (Flint et al., 2012; Porter and Martens, 2017). Host digestive enzymes of salivary, gastric, and pancreatic origin target the major classes of dietary nutrients, notably fat, protein, and cooked or non-resistant starches (Goodman, 2010; Iqbal and Hussain, 2009). In contrast, dietary fibers are degraded far less, if at all, by host enzymes and instead require members of the gut microbiota for transformation into host-absorbable short chain fatty acids (Macfarlane and Macfarlane, 2003). As a consequence, dietary carbohydrates play an important role in shaping the composition and physiology of the gut microbiota (David et al., 2014; Sonnenburg et al., 2016; Sonnenburg et al., 2010). Unlike the aforementioned nutrients, the digestive fates of nucleic acids (from diet, host or microbial origin), their component covalently linked ribo- and deoxyribonucleosides and cofactors or modifications built from similar molecules are less understood. In particular, the identities of common symbiotic human gut bacteria that are capable of utilizing these molecules and the most relevant source(s) and forms of these substrates remain obscure. Since enterohemorrhagic *E. coli* (EHEC) and other pathogenic *E. coli* have been shown to utilize nutrients like ribose or deoxyribose or associated nucleic acids/nucleosides (Fabich et al., 2008; Martinez-Jehanne et al., 2009), these substrates may represent unexplored nutrient niches that are competed for by commensal and pathogenic microorganisms and therefore help mediate colonization resistance against pathogens.

Some commensal and pathogenic human gut bacteria have known abilities to utilize free ribose or deoxyribose, as well as (deoxy)ribonucleosides and nucleic acids. Specifically, strains of mutualistic *Lactobacillus* (McLeod et al., 2011) and *Bifidobacterium* (Pokusaeva et al., 2010), as well as pathogenic and non-pathogenic *Escherichia coli* (Fabich et al., 2008) and *Salmonella enterica* (Harvey et al., 2011) have characterized mechanisms for ribose catabolism. Further, the ability of EHEC to prioritize ribose as a nutrient *in vivo* is thought to provide an advantage over commensal *E. coli* HS and may delineate different niches occupied by these strains (Maltby et al., 2013). Additional systems containing nucleoside-cleaving enzymes have been defined in *E. coli* and certain species of *Corynebacterium* isolated from feces (Hammer-Jespersen et al., 1971; Kim et al., 2006). One of the more interesting groups of nutrients in this category that can be used by some gut bacteria is DNA. As demonstrated in *E. coli,* DNA serves as a sole-carbon source through the action of competence genes and exonucleases (Finkel and Kolter, 2001; Palchevskiy and Finkel, 2009). However, mechanisms for assimilating exogenous RNA have not been explored.

Members of the phylum *Bacteroidetes* constitute a major portion of all bacteria found in the human gut, and species within this phylum devote large proportions of their genomes towards carbohydrate utilization via coordinately regulated polysaccharide utilization loci (PULs). A number of PULs have been investigated in great depth in model species like *Bacteroides thetaiotaomicron* and *Bacteroides ovatus* (Cuskin et al., 2015; Larsbrink et al., 2014; Luis et al., 2018; Ndeh et al., 2017). Most of the characterized systems target dietary or host polysaccharides, such as those found in plant cells, fermented foods or host mucosal polysaccharides, while others have been definitively linked to degradation of less common dietary substrates such as agarose and porphyran from edible seaweed (Hehemann, 2012; Pluvinage et al., 2018). Despite variations in the substrates they target, the cellular “Sus-like systems” encoded by Bacteroidetes PULs (Martens et al., 2009) are patterned in similar ways— each containing one or more TonB-dependent receptors (SusC homologs) and adjacently encoded substrate binding lipoproteins (SusD homologs). These two proteins form a complex with extensive protein-protein interactions (Glenwright et al., 2017) and work with a variable repertoire of cell surface and periplasmic carbohydrate-degrading enzymes, substrate binding proteins and regulators to bind, degrade and import their specific substrates. However, despite many studies examining the substrate specificity and function of Sus-like systems, many additional PULs have been identified in the genomes of gut and environmental *Bacteroidetes* without existing knowledge of their target substrates (Terrapon et al., 2018).

Here we describe a diverse set of gene clusters present in human gut and environmental *Bacteroidetes* that are regulated by ribose, but have likely evolved to target a variety of different ribose-containing nutrients. Using *Bacteroides thetaiotaomicron* (*Bt*) as a model, we investigated a PUL of unknown function that is upregulated *in vivo* in multiple dietary conditions, including in the absence of dietary fiber when this bacterium is forced to forage host-derived nutrients. This gene cluster contains two predicted ribokinases and a nucleoside hydrolase (NH) among other functions. Based on these predictions, we provide support for the hypothesis that this PUL targets ribose as a nutrient, which may in many cases need to be cleaved from covalently linked sources, and show that it confers a diet-specific competitive advantage to *Bt* during *in vivo* colonization. Surprisingly, the dietary condition during which this ribose utilization system (*rus*) is required is a high-fiber diet that supplies a variety of other carbohydrate nutrients. *Bt* does not require its two *rus*-encoded hydrolytic enzymes, or SusC/D-based transport for this diet-specific competition. Instead, ribokinase function is essential, suggesting that sensing and metabolism of simpler ribose and ribose-phosphate derivatives provides the key metabolic advantage in this particular dietary condition. Taken together, our results reveal that a variety of host-associated and terrestrial bacteria, including numerous human gut symbionts have evolved mechanisms to scavenge ribose from various sources. The common regulation of a family of highly diversified PULs by a ubiquitous simple sugar that occurs in a variety of molecular contexts, such as nucleic acids, co-factors, modifications (ADP- and poly-ADP-ribose) and bacterial capsules, suggests that these systems have been adapted at the enzymatic level to release ribose from these varied sources, allowing Bacteroidetes to expand their colonization of diverse nutrient niches.

## Results

### A ribose-inducible gene cluster is highly active *in vivo* and required for fitness in a diet-dependent fashion

Members of the prominent human gut bacterial phylum *Bacteroidetes* typically encode coordinated degradative functions within discrete gene clusters called polysaccharide utilization loci (PULs), facilitating identification of functions that work together to access a particular nutrient (Martens et al., 2009). Previous work using gnotobiotic mice in which *B. thetaiotaomicron* (*Bt*) is the only colonizing bacterium identified a locus (*BT2803-2809*) for which all individual genes are upregulated between 10- and 139-fold *in vivo*, including in mice fed diets with high and low dietary fiber (**Fig. 1A**). Under low fiber conditions, *Bt’s* physiology shifts to expression of genes involved in host glycan foraging (Bjursell et al., 2006; Martens et al., 2008; Sonnenburg et al., 2005). Thus, the corresponding expression of *BT2803-09* in the absence of dietary fiber suggests that it also targets endogenous nutrients. Interestingly, the genes in this locus were also most highly expressed in neonatal mice still consuming mothers milk, which is not only fiber-deficient but also contains milk-derived ribose and nucleosides (Schlimme et al., 2000). Typically, PULs involved in host glycan foraging encode enzymes required for liberating nutrients from mucus glycoproteins: fucosidase, sulfatase, β-galactosidase and β-N-acetylhexosaminidase (Bjursell et al., 2006; Sonnenburg et al., 2005). However, some predicted enzymes encoded in the *BT2803-09* PUL (nucleoside hydrolase and ribokinases) suggest a role in harvesting ribose from substrate(s) such as nucleosides or RNA, which could be of host or gut bacterial origin. Previous studies have determined that *Bt* can grow on ribose as a sole carbon source (Martens et al., 2011). However, the genes involved, the relevant source(s) of this sugar, and whether it involves enzymatic liberation from complex sources remain unknown.

**Figure 1.**
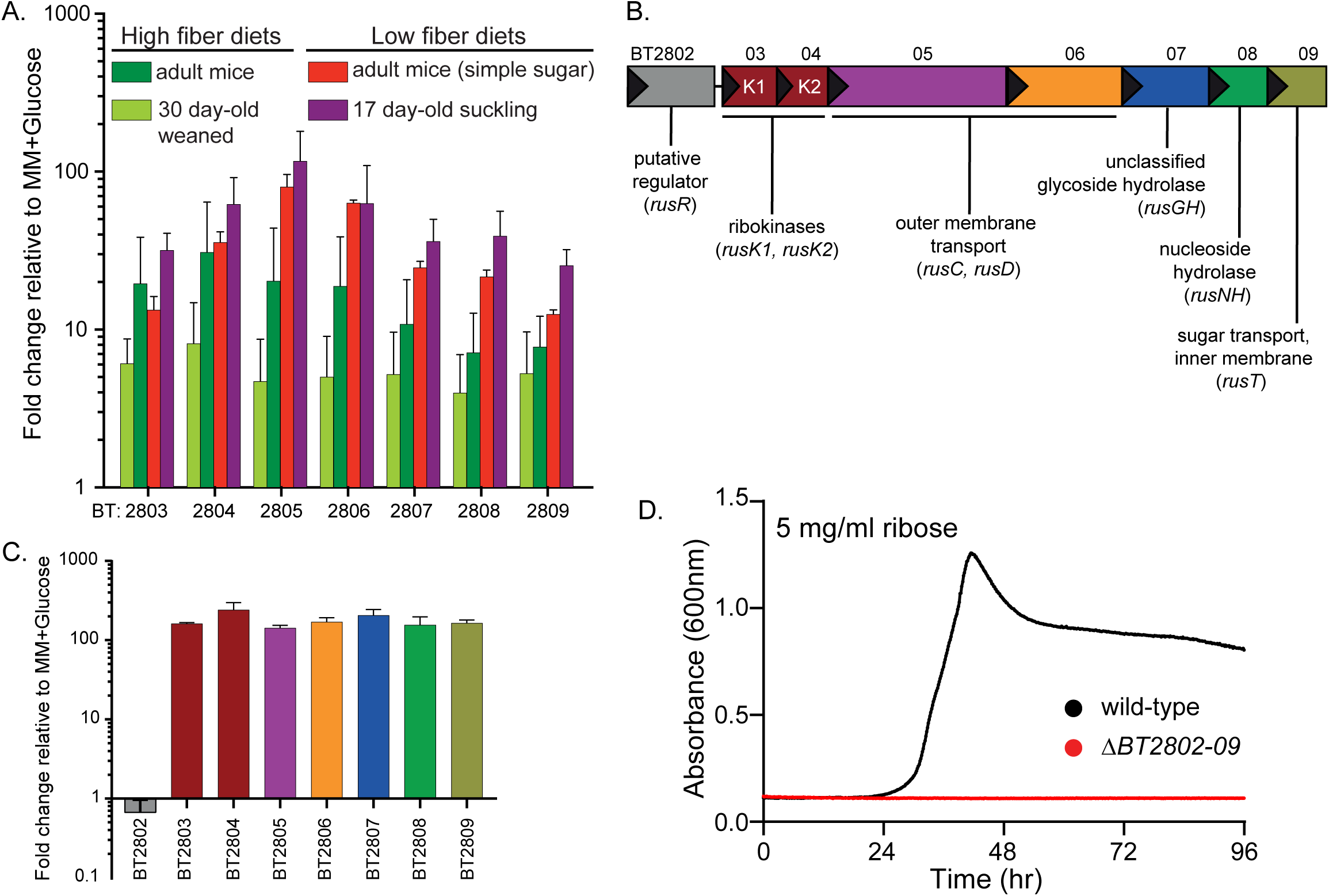
*Bt* upregulates a PUL for ribose metabolism *in vivo* and *in vitro* in response to ribose. (A) *In vivo* gene chip data (Bjursell et al., 2006; Sonnenburg et al., 2005) displaying fold-change relative to *in vitro* growth in minimal medium (MM), plus glucose for *BT2803-09* in mice fed high fiber (dark and light green bars) or low fiber diets, including pre-weaned, suckling mice (red and purple bars, respectively). (B) Genomic architecture of the *rus* locus spanning genes *BT2802-09* with names and predicted functions. (C) *In vitro* transcriptional response of *rus* genes in *Bt* grown on ribose as a sole carbon source, measuring fold-change relative to growth on a MM+glucose reference (n=3, error bars are SD of the mean). (D) Growth on minimal media containing ribose (5 mg/ml) as the sole carbon source in wild-type *Bt* (black) or a strain lacking *rus* (red) through 96 hours of incremental readings (minimum of n=5 separate replicates).

The architecture of the PUL spanning *BT2803-09* revealed several unique features compared to other PULs activated in low fiber diets (**Fig. 1B)**. The immediate upstream gene (*BT2802*) is predicted to have DNA-binding motifs and may act as a regulator but shares no homology to regulators previously associated with PULs, the next two genes are predicted ribokinases (*BT2803* and *BT2804*), followed by genes encoding homologs of the *Bacteroides* SusC and SusD outer-membrane proteins (*BT2805, BT2806*), a glycoside hydrolase of unassigned family and function (*BT2807*), a predicted nucleoside hydrolase (*BT2808*), and a sugar permease (*BT2809*). The enzymes encoded in this PUL suggested the hypothesis that it is responsible for *Bt’s* ability to catabolize ribose and possibly liberate it from more complex sources, such as RNA, nucleosides, or cofactors. To test if this gene cluster is transcriptionally responsive to growth on ribose, we performed *in vitro* growth experiments in minimal-medium (MM) containing ribose as the sole carbon source and measured expression of the genes *BT2803-09*. All genes were activated 142-240 fold during growth on ribose compared to growth on a MM-glucose reference (**Fig. 1C**). We next examined the contribution of this PUL to ribose catabolism by deleting the entire PUL and upstream gene (*BT2802*). Consistent with an essential role in ribose catabolism, loss of the PUL eliminated the ability to grow on free ribose (**Fig. 1D**). Based on these findings, we classified this PUL as the *Bt* ribose utilization system, *rus*, with gene annotations *rusR* (putative regulator), *rusK1* and *rusK2* (ribokinases 1 and 2), *rusC* (SusC-like), *rusD* (SusD-like), *rusGH* (glycoside hydrolase), *rusNH* (nucleoside hydrolase), and *rusT* (transporter), for genes *BT2802-09* respectively. These *in vitro* results, combined with the observation that *rus* exhibits high activity in the gnotobiotic mouse gut, led us to hypothesize that the ability to utilize endogenous sources of ribose-containing nutrients is advantageous *in vivo* during fiber-deficient diets.

To test the above hypothesis, we inoculated 6-8 week old, germfree (GF) female Swiss-Webster mice with a mixture of wild-type and Δ*rus Bt* strains (∼10^8^ total cfu/mouse, equal amounts of each) and maintained mice on either a fiber-rich (FR) diet containing several unprocessed plant-derived fiber polysaccharides or a fermentable fiber-free (FF) diet consisting mainly of glucose, protein, lipids, and cellulose (Desai et al., 2016). We measured the relative abundance (by qPCR) of each strain for 42 days in DNA extracted from feces. Surprisingly, and in contrast to our initial hypothesis, the Δ*rus* strain was strongly outcompeted (∼100-fold) only in mice fed the FR diet (**Fig. 2A**). In contrast, in mice fed the FF diet, Δ*rus* exhibited similar abundance to wild-type throughout the experiment (**Fig. 2B**). A similar competitive defect of the Δ*rus* strain in mice fed the FR diet was observed in separate experiments with 12-week-old female mice and 6-8 week old male mice (**Fig. S1A**,**B**), suggesting the competition is not influenced by sex or age within the range tested. The defect associated with the FR diet was not due to lack of colonization or persistence *in vivo,* as the levels of each strain were similar over time in mice colonized with either strain alone (**Fig. S1C**,**D**). Additionally, the defect seen in the FR diet could not be attributed to the wild-type strain exhibiting different expression of the *rus* PUL, as wild-type *Bt* exhibited similarly high levels of *rus* expression in mice fed either diet when present alone or in competition with the Δ*rus* mutant (**Fig. S1E**). GC-MS analysis of the two diets revealed that ribose was present only in the FR diet, in levels similar to other common monosaccharides, and in an acid-hydrolyzable (*i.e*., covalently linked) form, but not detectably as a free sugar. This suggested the presence of a ribose-containing molecule(s), such as RNA, nucleosides or cofactors, which may be scavenged by *Bt* in mice fed this diet (**Fig. S1F**). However, in cecal contents of mice monoassociated with wild-type or Δ*rus* strains and fed the FR diet, ribose was undetectable while other sugars present in acid-hydrolyzed extracts of the uneaten FR diet—and likely deriving from fibers such as arabinan and arabinoxylan—could still be measured (**Fig. S2**, note that we determined through control experiments that the limit of detection of ribose in the complex milieu of cecal contents is near the amount observed in the FR diet, although the levels of other sugars seems to have been concentrated as much as 2-fold during digestion). This unexpected result led us to conclude that although ribose is present in the FR diet it may be depleted or absorbed during transit through the upper GI, such that it is not detectable even in the cecal contents of mice colonized by Δ*rus*, which cannot use this sugar. Nevertheless, to directly test if dietary ribose from different sources can impact *Bt* in the distal gut, we colonized three separate groups of GF mice with a mixture of wild-type and Δ*rus* strains and maintained them on the FF diet that does not elicit a competitive defect for Δ*rus*. After 14 days of stable competition between strains, water was supplemented with either 1% ribose, 1% RNA, or 1% nucleosides (0.25% w/v each of uridine, thymidine, 5-methyl uridine, and cytidine). The results clearly show that free ribose present in the drinking water elicits a competitive fitness defect for the Δ*rus* strain similar in magnitude and with slightly faster timing, to the defect in mice fed the FR diet (**Fig. 2C)**. In contrast, little defect, if any, was observed in mice switched to water containing the nucleoside mix or RNA (**Fig. 2D**,**E**). Similar to mice fed just FR or FF, there was similar expression of the *rus* locus in all of the supplemented water conditions, suggesting that levels of *rus* expression did not account for the various fitness outcomes (**Fig. 2F**). While our findings above imply that little free ribose is present in the FR diet and even the acid-hydrolyzed, covalently-linked form(s) may be removed before reaching the distal gut, our results with dietary additions of ribose-containing nutrients clearly indicate that free dietary ribose, but not RNA or nucleosides, is a form of this nutrient capable of driving abundance changes in *Bt* populations.

**Figure 2.**
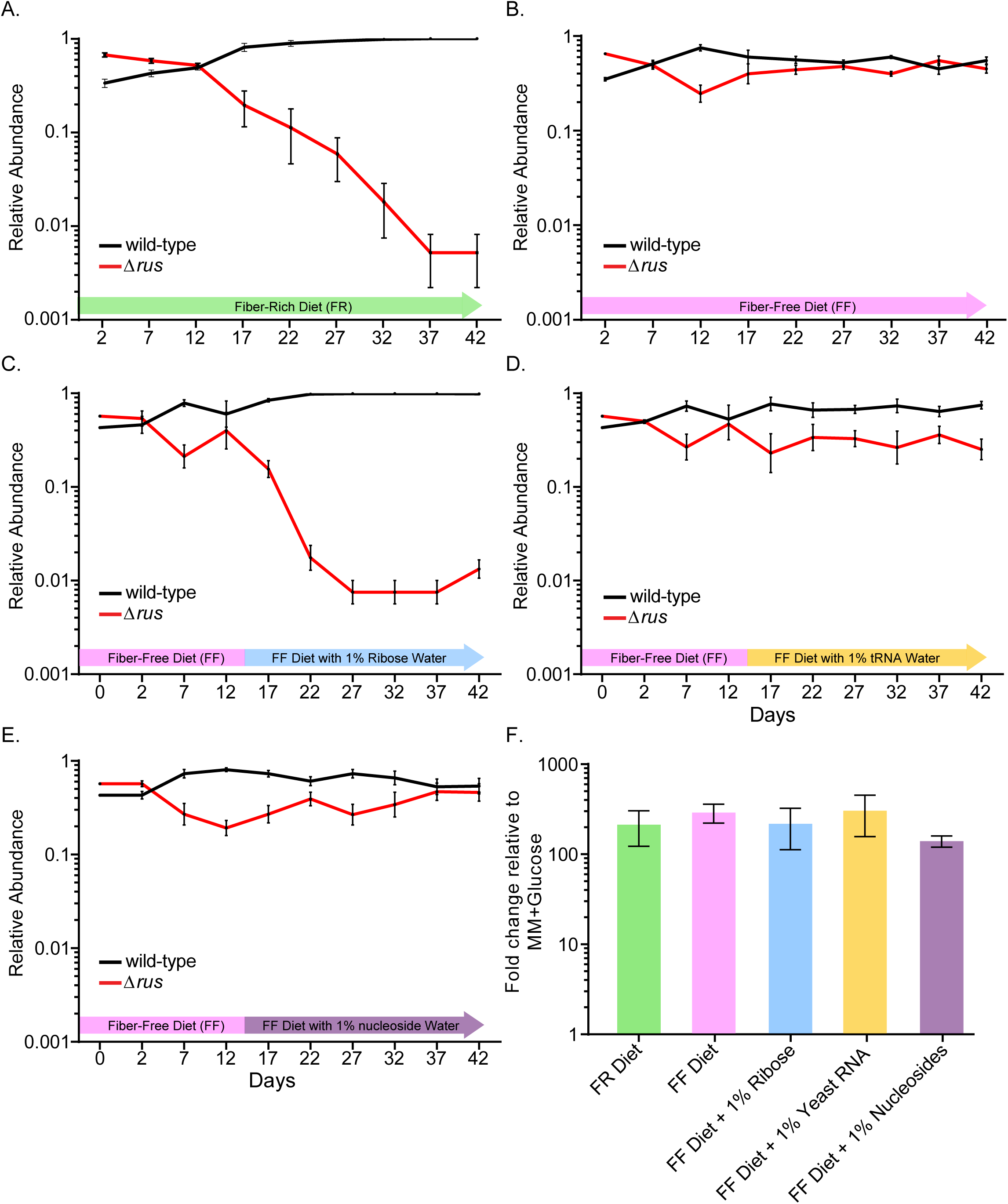
The locus *BT2802-09* confers a competitive advantage *in vivo* in a diet-dependent context. For all panels, relative abundance is shown on a log scale for wild-type (black) and Δ*rus, BT2802-09* (red) strains and as measured by qPCR from fecal samples of gnotobiotic mice (n=4 for each experiment) colonized with a mixture of the two strains indicated at day 0. (A) Mice continuously fed a high fiber diet (green arrow) for the entire duration. (B) Mice continuously fed a low fiber diet (pink arrow) for the entire duration. (C-E) Mice initially maintained on the FF-diet for two weeks and then at day 14 given *ad libitum* access to water containing 1% w/v ribose (C), 1% w/v of RNA from type IV Torula yeast RNA (D), or a 1% w/v mixture of nucleosides (0.25% each of uridine, cytidine, thymidine, and 5-methyl uridine). In each panel the duration of the water supplementation is shaded in either blue, orange, or purple, representing the different substrates. (F) *rus* transcript levels measured by qRT-PCR probing the *rusC* gene from *in vivo* cecal contents, bar colors correspond to the background shading in panels A-E. Error bars in all panels display the standard error of the mean for each time point.

### A subset of ribose-utilization functions is required for competitive colonization in mice

The experiments described so far utilized a mutant lacking all 8 *rus* genes, but only a subset of the functions may be important for competition with wild-type in mice fed the FR diet. Because biochemical approaches failed to reveal a clear ribose source that drives the competitive advantage associated with *rus* expression in the FR diet, we took a molecular genetic approach to probe the required enzymatic and transport functions. We constructed single and double gene deletions based on predicted functionality (**Fig. 1B**), performing additional competitive colonization experiments in FR diet-fed mice. Each individual mouse group was inoculated with wild-type *Bt* and one of the following competing strains (Δ*rusK1/2,* Δ*rusC/D,* Δ*rusGH/NH,* Δ*rusT,* or Δ*rusR*). Surprisingly, the only strain that exhibited a competitive fitness defect similar to the full Δ*rus* mutant was Δ*rusK1/K2*, which lacks both predicted ribokinases (**Fig. 3A**). In contrast, the other four deletion strains exhibited equal or better competition compared to wild-type (**Fig. 3B-E**). Notably, the Δ*rusGH/NH* strain, which lacks *rus*-associated hydrolase functions, exhibited a significant competitive advantage (∼100-fold better than wild-type). These results clearly suggest that the required functions underlying the competitive defect in the Δ*rus* strain are encoded by the *rusK1* or *rusK2* genes, while expression of the other genes provide no advantage in this context, and perhaps even a fitness disadvantage in the FR diet. To further address which of the ribokinases is important *in vivo* we repeated the above competition with single Δ*rusK1* and Δ*rusK2* deletion strains. Each of these single kinase mutants also competed better than wild-type, suggesting that they are redundant and need to be lost together to elicit a defect (**Fig. 3F**,**G**). As in previous experiments, we could not attribute variations in competitive behavior to a significant difference in *rus* expression in wild-type *Bt*, which was elevated in all cases (**Fig. 3H**).

**Figure 3.**
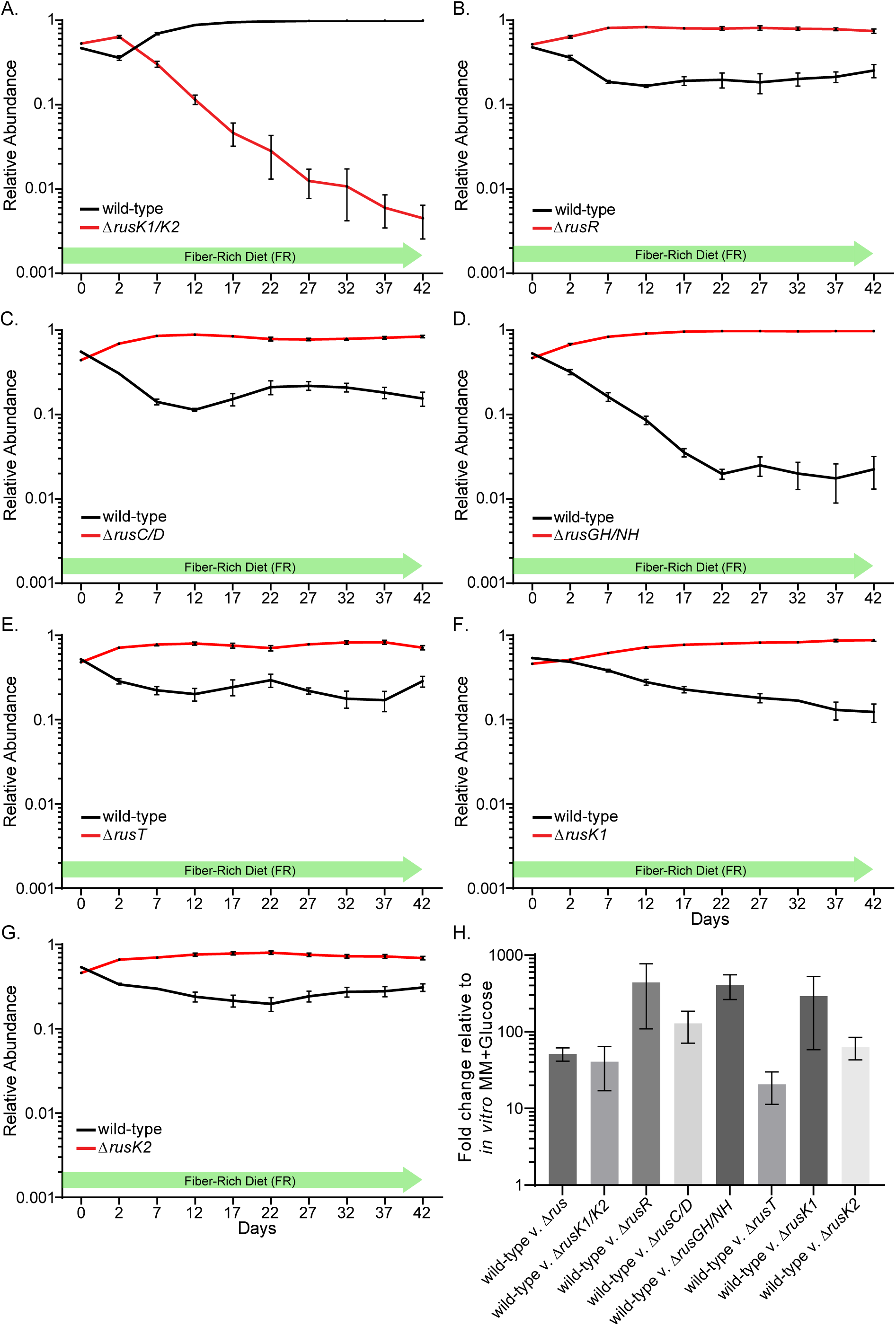
Ribokinases *rusK1*/*K2* are sufficient to confer a competitive advantage *in vivo.* (A-G) In all experiments, 6-8 week old germfree Swiss-Webster mice were fed the fiber rich (FR) diet and gavaged with a mixture of both wild-type *Bt* (black) and the mutant strain indicated (red) in nearly equivalent levels. Relative abundance is displayed over a 42-day experiment identically to Figure 2. In all panels error bars represent standard error of the mean of four biological replicates. (H) *rus* transcript levels measured by qRT-PCR probing the *rusC* gene from *in vivo* cecal contents for bacterial populations in each panel.

### A subset of Rus functions is required for sensing and utilization of ribose containing nutrients *in vitro*

The results described above clearly indicate a diet-specific advantage for *Bt* strains that possess the *rus*-encoded ribokinases. To further define this system’s function, we tested our panel of deletion mutants in a variety of growth conditions, including free ribose, nucleosides, RNA, and other sources of this sugar. Consistent with our *in vivo* data, a mutant lacking both *rusK1* and *rusK*2 could not grow on free ribose (**Fig. 4A**). However, arguing against purely redundant functions as concluded above, the mutant lacking just *rusK2* displayed a complete loss of growth phenotype, while a mutant lacking only *rusK1* reproducibly displayed a substantial growth lag, but eventually grew with slightly slower rate compared than wild-type (**Fig. 4B**,**C**). The delayed growth phenotype of this mutant might be attributed to genetic suppressor mutations or another heritable alteration, since cells that eventually grew were able to grow quickly on ribose after being isolated and passaged in rich media (**Fig. S3A**). Deletion of the flanking gene *rusR*, a candidate transcriptional regulator, was also unable to grow on ribose, suggesting that, although it is not transcriptionally activated in response to ribose, its product plays an essential role in ribose catabolism (**Fig. 4D**). The Δ*rusT* strain exhibited an increased lag, slower growth rate and lower overall growth level compared to wild-type (**Fig. 4E**). Unlike the Δ*rusK1* mutant this mutant did not exhibit increased growth after passaging and re-testing, suggesting that suppressor mutations are not involved and rather another, lower-affinity pentose sugar permease is present to import ribose. Lastly, the Δ*rusC/D* strain consistently exhibited a longer growth lag on ribose, suggesting that although ribose should freely diffuse across the outer membrane, the RusC/D complex on the cell surface might increase affinity for this sugar leading to accelerated growth (**Fig. 4F**, reference growth of wild-type *Bt* on ribose is shown here). All of the other single or double deletion mutants tested (Δ*rusC,* Δ*rusD,* Δ*rusGH,* Δ*rusNH,* Δ*rusGH/NH*), exhibited no measurable difference in growth parameters compared to wild-type *Bt* when grown on ribose (**Fig. S3B-F**, **Table S1**).

**Figure 4.**
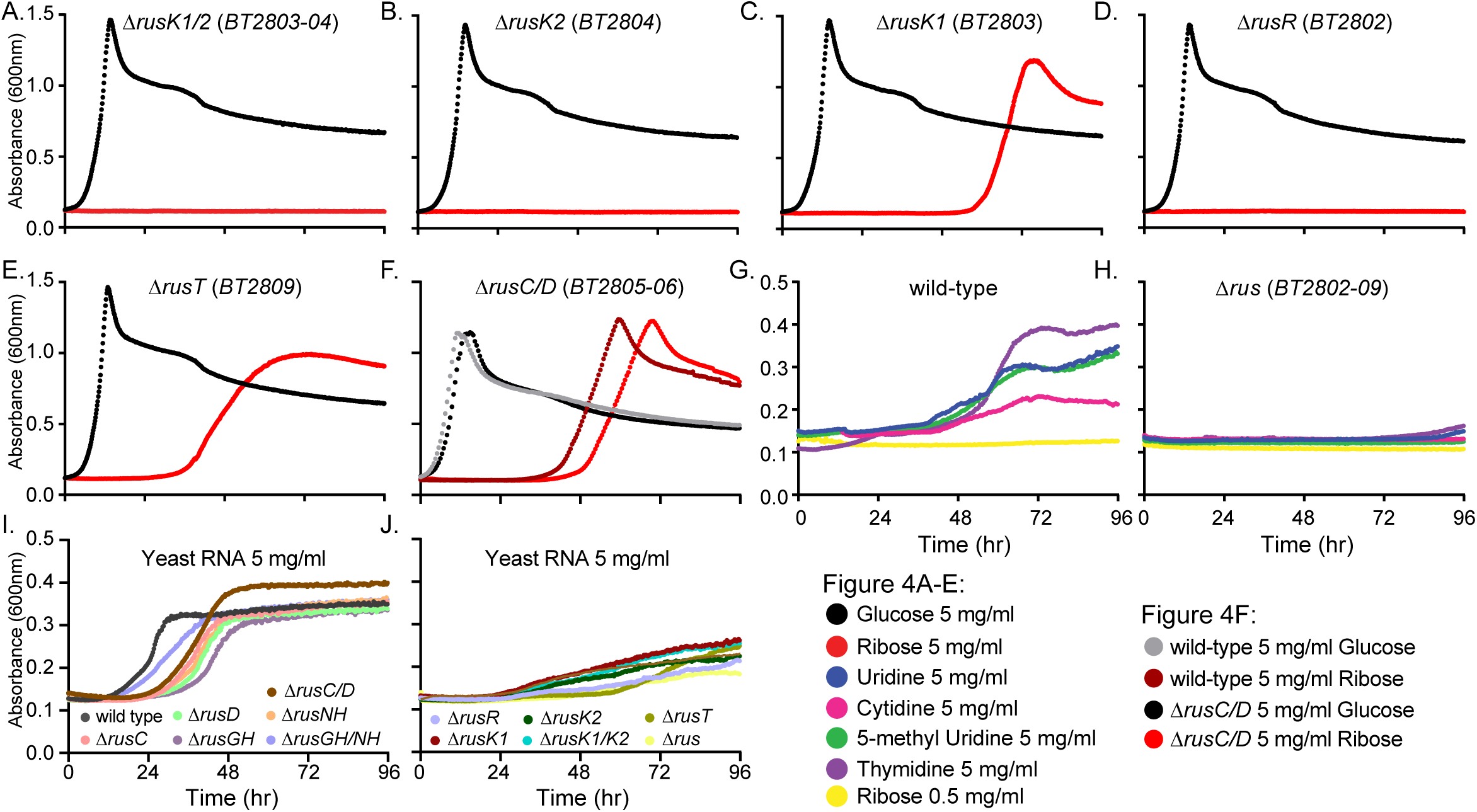
The *Bt rus* PUL encodes functions required for growth on ribose containing nutrients. (A-E) Growth curves of *rus* deletion strains on ribose (red) or glucose (black) that exhibit defects for growth on ribose compared to wild-type *Bt* (see panel F. for a wild-type reference curve for growth on ribose). (F) Growth of wild-type and the Δ*rusC/D* strains are shown for glucose and ribose, revealing slightly weaker growth for the *rusC/D* mutant. (G-H) Growth of wild-type (G) or Δ*rus* (H) on different nucleosides in the presence of 0.5 mg/ml ribose (yellow line is media with only 0.5 mg/ml ribose). (I-J) Growth of wild-type and *rus* deletion strains on 5 mg/ml yeast RNA and media containing RNase A and IAP. Strains displaying similar growth profiles as wild-type are in (I), while strains showing a reduction in growth are shown in (J). Note that in panels G-H the y-axis scale is reduced. Curves shown in each panel are the average of a minimum of 8 technical replicates.

Since they are larger and more complex, we hypothesized that utilization of molecules in which ribose is covalently linked to other ligands would require additional *rus*-encoded functions. To test this, we assayed growth of our *rus* mutants and wild-type *Bt* on nucleosides and RNA. Wild-type *Bt* displayed poor growth, when observed at all, on all nucleosides tested (uridine, cytidine, 5-methyl uridine, thymidine, inosine, xanthosine, adenosine) as well as on RNA (**Fig. S3G**,**H**, **Table S1**). We hypothesized that free ribose may need to be present to activate transcription of the *rus* locus, generating proteins necessary for catabolism of these substrates. To test this, we determined a concentration (0.5 mg/ml) at which ribose elicited strong *rus* expression but little if any measurable growth (**Fig. S3I**,**J**). We then re-evaluated the ability of wild-type *Bt* to grow on nucleosides in the presence of this low ribose concentration, observing considerably higher levels of total growth on pyrimidine nucleosides (**Fig. 4G**). While growth was still comparably poor relative to growth on pure ribose, increased growth was not observed when we doubled the nucleoside concentrations, suggesting that something else about growth on nucleosides limits growth (**Fig. S3K**). Growth on nucleosides was eliminated in mutants lacking the full locus (**Fig. 4H**), either or both ribokinases (*rusK1, rusK2*, and *rusK1/K2*), the candidate regulator (*rusR*) and the putative transporter (*rusT*). Each of these phenotypes was similar to those observed for growth on free ribose, except the Δ*rusK1* and Δ*rusT* mutants, which eventually grew with reduced rate on ribose, but not on nucleosides (**Fig. S3L-P**). Growth on RNA alone was not observed, even after addition of ribose, suggesting that *Bt* does not produce sufficient extracellular RNAse and phosphatase enzymes required to liberate nucleosides from this polymer. Therefore, we tested if exogenous RNase A and intestinal alkaline phosphatase (IAP), which are present in the gut from host pancreatic secretions (RNAse) or native to the enterocyte brush boarder and secreted in luminal vesicles (IAP), could enhance growth on RNA at physiologically relevant concentrations (McConnell et al., 2009; Weickmann et al., 1984). When these host-derived enzymes were supplemented in media, growth on RNA was appreciably greater than in their absence (**Fig. 4I**), which was not attributable to *Bt* growing on the exogenous enzymes alone (**Fig. S3Q**). As with individual nucleosides, reductions or eliminations in growth on enzyme-degraded RNA were observed in mutants lacking the entire *rus* locus, *rusK1, rusK2, rusK1/K2, rusT*, and *rusR* (**Fig. 4J**). In addition to free nucleosides and those derived from RNA, we also determined that *Bt* is able to utilize the pentoses deoxyribose and lyxose, as well as ADP-ribose and UDP-galactose: each of these required the presence of both a low amount of ribose and the *rus* locus. Sixteen additional substrates did not support *Bt* growth under any conditions (**Table S1**).

Inconsistent with our initial hypothesis, mutants lacking functional *rusC, rusD, rusGH, rusNH, rusC/D* or *rusGH/NH* encoded products, exhibited total growth levels comparable to wild-type on both nucleosides and degraded RNA (**Fig. 4I**, **S3R-W**). These results suggested that other genes in *Bt* encode functions responsible for cleavage and utilization of free nucleosides and those liberated from RNA. To identify other functions involved in utilization of nucleosides, we searched the *Bt* genome for functions from known nucleoside scavenging systems (NSSs), identifying several candidates. We made deletions of 4 genes predicted to encode nucleoside cleavage and import functions, *BT0184, BT1881, BT4330,* and *BT4554,* which are predicted to encode a uridine kinase, a purine nucleoside phosphorylase, a nucleoside permease, and a pyrimidine nucleoside phosphorylase, respectively. We tested growth of these strains on nucleosides (**Fig. 5A-D**) and only one strain (Δ*BT4554*) displayed loss of growth on all nucleosides tested, suggesting that it encodes an essential enzyme for cleaving nucleosides and works upstream of also required *rus* functions. The Δ*BT4330* mutant also exhibited reductions in growth on uridine, cytidine, and 5-methyl uridine, while only slight defects were seen in thymidine (**Fig. 5A-D**). Further, the Δ*BT0184* mutant displayed enhanced growth that began quicker than wild type and reached a higher total growth level on all nucleosides, except the deoxyribonucleoside thymidine. This phenotype could be due to its role in 5’ phosphorylating scavenged nucleosides and shunting them towards anabolic pathways, such that when it is deleted catabolic growth is enhanced. Lastly, Δ*BT1881* did not display any detectable growth defects compared to wild-type, suggesting that the product of this gene is not essential for pyrimidine nucleoside catabolism. Interestingly, although growth on nucleosides in some NSS mutants were reduced or eliminated compared to wild-type, this phenotype did not extend to growth on RNA, as the mutant strains exhibited similar levels of growth as wild-type (**Fig. S4A**,**B**). These results suggest that, while Rus functions are required to use RNA-derived nucleosides, the NSS functions interrogated here are not essential for catabolism of RNA-derived nucleosides, which might vary in their nucleoside ratios or oligomer length and could be assimilated via additional pathways.

**Figure 5.**
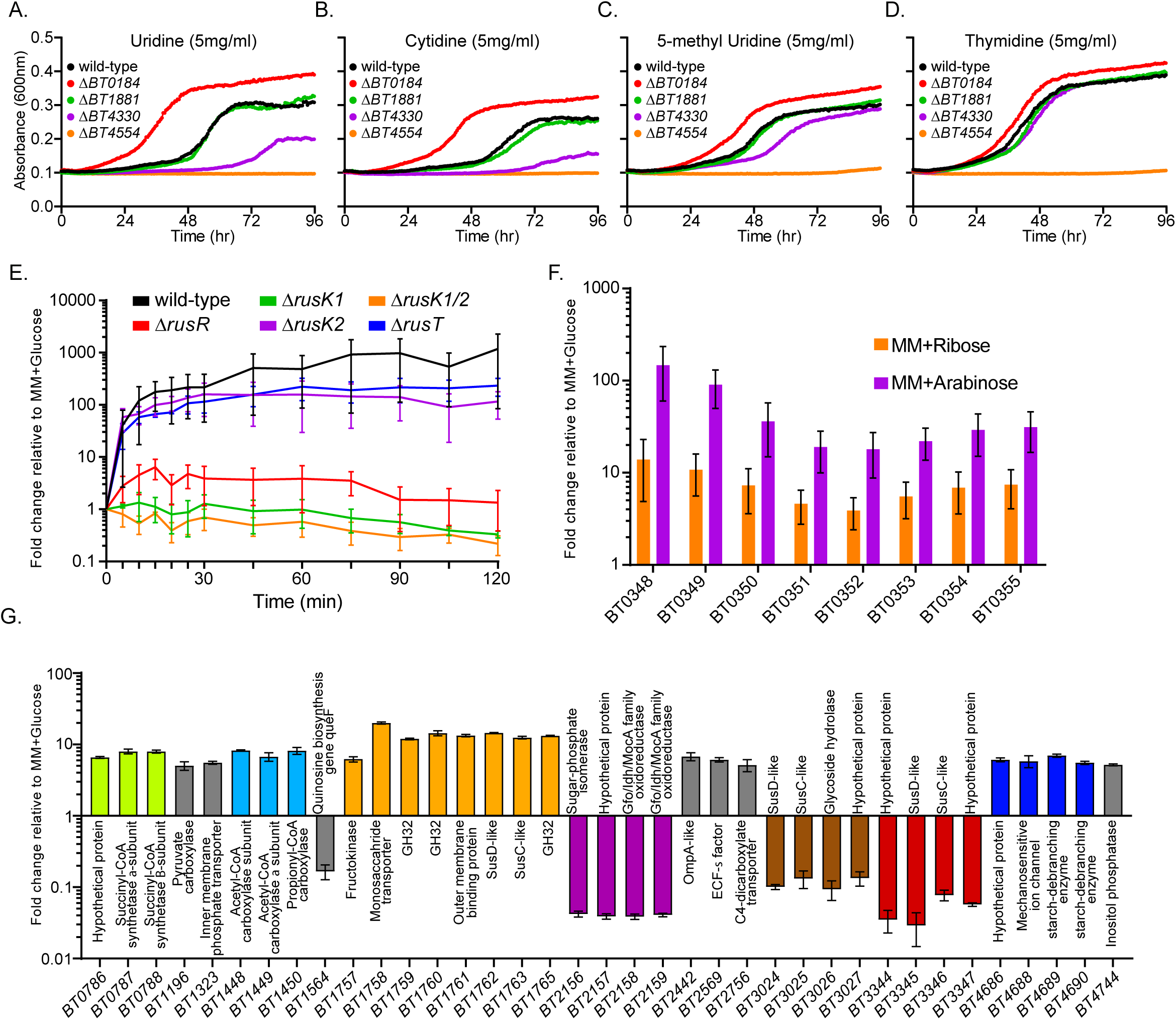
Requirement of some nucleoside scavenging systems, dynamics of *Bt* rus activation and global responses to ribose. (A-D) Growth curves of predicted nucleoside scavenging mutants (Δ*BT0184*, red; Δ*BT1881*, green; Δ*BT4330*, purple; or Δ*BT4554*, orange) and wild type *Bt* (black) on uridine, cytidine, 5-methyl uridine or thymidine. (E) *Bt rusC* transcriptional response as measured by qRT-PCR over time after mid-log phase cells actively growing in MM-glucose (5 mg/ml) were transferred to MM-ribose (5 mg/ml). Samples were taken every 5 minutes for the first 30 minutes and then every 15 minutes after until 120 minutes. Strains are color coded according to the legend: wild type (black), Δ*rusK1* (green), Δ*rusK1/K2* (orange), Δ*rusR* (red), Δ*rusK2* (purple), and Δ*rusT* (blue). Shown are the averages of n=3 separate experiments (different days), error bars indicate the standard error of the mean. (F) Transcriptional responses of genes in the arabinose gene cluster *BT0348-0355* during growth on 5 mg/ml ribose (orange) or 5 mg/ml arabinose (purple), compared to *Bt* grown in MM-glucose (n=3 bars and error bars represent mean plus standard error). (G) Select RNAseq results indicating a global transcriptional response during growth on MM-ribose compared to a MM-glucose reference by fold-change in gene expression in each condition. Matching bar colors indicate genes are in the same locus or PUL. Additional genes listed in gray bars are not genomically linked to adjacent-coordinately regulated genes. Bars represent the mean plus standard deviation of 3 replicate experiments.

### Rus enzymes are active towards ribose-containing substrates or nucleosides

The results described above suggest that the product of the *rusNH* gene, if functional, is superfluous to pyrimidine nucleoside salvage, since deletion of *BT4554* eliminated growth on these nutrients. To test if this enzyme is actually functional, we produced a recombinant *Bt* RusNH by over-expression in *E. coli* and performed substrate cleavage assays. We first used *p*-nitrophenyl-β-D-ribofuranoside (pNP-ribose), on which this enzyme was active and determined the pH optimum to be 6.7. We next tested the cleavage specificities and affinities of ribo- and deoxyribonucleosides in a UV-based assay (Liang et al., 2008), observing that RusNH has broad, but relatively weak activity compared to other nucleoside hydrolases towards all nucleosides tested (**Table S2**). This broad range of catalytic activities suggests that this enzyme belongs to the inosine-uridine-preferring family of nucleoside hydrolases (IUNH) as predicted by annotation. However, despite containing the canonical N-terminal DXDXXXDD motif involved in binding of ribose and coordination of Ca^2+^ ions (**Fig. S4C**), the kinetic values of RusNH are not within the range of previously characterized IUNH hydrolases from other organisms (**Table S2**) (Parkin et al., 1991; Shi et al., 1999; Versées and Steyaert, 2003). Although we attempted to directly measure enzyme affinities by km determination, the activity was too weak to reach Vmax at the concentrations tested, further suggesting that RusNH is not primarily responsible for nucleoside cleavage.

Because RusNH has relatively weak activity towards nucleosides, we hypothesized that the predicted glycoside hydrolase, RusGH, could have activity on nucleosides since it has not been assigned a previously defined glycoside hydrolase (GH) family. A potentially important role for RusGH was further suggested by its possible location on the cell surface, which was suggested by a signal peptidase II secretion-lipidation signal and confirmed by antibody staining, whereas RusNH lacks this signal and appears to be secreted into the periplasm (**Fig. S4D**, data not shown for RusNH, which was not similarly detected on the cell surface despite being detectable in whole cell lysates by western blot). We therefore produced recombinant RusGH and tested a broad-range of pNP-based substrates in several buffer conditions and found optimum conditions to be pNP-ribose at pH 9.0 (**Table S2**). Arguing against a major role in cleavage of any of the substrates tested, RusGH displayed only weak activity on pNP-ribose that was too slow for accurate kinetic determinations and no detectable activity on other pNP substrates after 24 hours. Interestingly, the weak activity displayed was calcium or divalent cation dependent as addition of EDTA completely eliminated activity (**Table S2**). When RusGH was tested for the ability to cleave nucleosides for 24h, no liberation of ribose was observed. Additional testing on a panel of glycans that are capable of supporting *Bt* growth failed to reveal any additional activity. Thus, although the *Bt* Rus harbors two enzymes with demonstrable but weak activities, roles for these enzymes remains enigmatic, although it is possible that larger polymers exist that are the targets for these enzymes.

### Dynamics of *rus* transcript activation and global responses to ribose catabolism

Our *in vivo* and *in vitro* data support an important role for some Rus functions in utilization of ribose and ribonucleosides, although a critical part of the latter pathway hinges on the function of an unlinked gene, *BT4554*. Because Rus function and a small amount of free ribose is essential for utilization of nucleosides via the BT4554 phosphorylase, we sought to determine the requirements for activating expression of *rus* genes as well as the presence of other global responses that ribose may induce. We hypothesized that the critical Rus functions for responding to free ribose are the reaction products from one or both kinases, RusK1, RusK2, and also require the putative regulator RusR, and the permease, RusT. To test this, we examined the kinetics of *rus* transcriptional responses when *Bt* was exposed to ribose. We grew our wild-type and mutant strains in medium containing glucose as a sole carbon source, washed them in a carbohydrate-free medium, transferred the bacteria into medium containing ribose as the only carbon source and monitored *rus* transcript accumulation over time (**Fig. 5E**). Our results show that wild-type *Bt* achieves close to maximum activation between 15-30 minutes post-exposure, with continued slow increase in expression. Interestingly, the Δ*rusK2* strain, which cannot grow on ribose, still generated near wild-type levels of transcript on the same time scale. In contrast, the Δ*rusK1* mutant, which exhibits an extensive lag before growth on ribose, was unable to quickly generate transcript on a 2 hr time scale (**Fig. 5E**), but eventually achieves near wild-type *rus* expression once actively growing on ribose (**Fig. S4E**). This latter result is consistent with our previous observation with the *Bt* starch utilization system (Cameron et al., 2014) that growth phenotypes characterized by protracted lag periods are in some cases due to deficient ability to sense an activating sugar cue. As expected based on the single kinase deletions, the Δ*rusK1/K2* double mutant was unable to generate transcript, while the Δ*rusT* exhibited only slightly lower expression than wild-type (**Fig. 5E**). The Δ*rusR* mutant only achieved partial (∼10%) activation in response to ribose, which supports the idea that RusR is required for positive transcription activation and the partial expression could be due to the absence of glucose repression during ribose exposure. Finally, we measured *rus* expression dynamics in our Δ*rusC* and Δ*rusD* strains, with the hypothesis that these outer membrane proteins may increase the cell’s affinity for ribose leading to more rapid activation, but failed to detect any differences in expression compared to wild type. (**Fig. S4F**). Further, to rule out nucleosides (processed to ribose-1-P via BT4554) serving as a possible inducing molecule we monitored *rus* transcript over time when wild type *Bt* was exposed to either uridine or inosine in the absence of ribose and did not see *rus* activation (**Fig. S4G**).

The two ribose-inducible kinases encoded in the *Bt rus* locus are predicted to generate products that are part of the pentose phosphate pathway (PPP), with known ribokinases adding phosphate to the 5’ position, although we cannot rule out generation of (1-, 5- or 1-/5-phosphate). Thus, we hypothesized that growth on exogenous ribose may affect expression of a more global regulon that could contribute to the *in vivo* competitive defect associated with the FR diet. We initially probed expression of genes involved in metabolizing the other pentoses, xylose and arabinose, revealing that growth on ribose leads to increased expression of genes involved in arabinose utilization, but only to ∼10-20% of levels when grown directly in arabinose (**Fig. 5F**), however the same effect was not seen for genes involved in xylose metabolism (**Fig. S4H**). To extend these results, we performed RNAseq-based whole-genome transcriptional profiling on wild-type *Bt* grown on ribose or glucose to address if growth on ribose elicits a broader metabolic response. The data indeed reveal a global response in which 81 genes are differentially expressed according to the parameters used. Unexpectedly, many of the genes (46%) belong to other PULs or metabolic pathways, with most of the remaining genes encoding hypothetical functions or undefined pathways. Notable changes included upregulation of a previously defined PUL for fructose and β2,6-linked fructan metabolism (*BT1757-1765;* average upregulation of 15-fold), which interestingly liberates fructose that initiates the PPP (Sonnenburg et al., 2010). At the same time, two other PULs of unknown specificity were repressed (*BT3024-3027, BT3344-3347*). Further, *Bt* has a partial TCA cycle pathway (Pan and Imlay, 2001) of which several genes leading to generation of succinate and propionate were upregulated, while genes predicted to participate in sugar-phosphate isomerization and metabolism, are strongly repressed (*BT2156-2159;* average of 24-fold) (**Fig. 5G**). An experiment to test the hypothesis that cross-regulation between ribose metabolism and the fructan PUL is the cause of the FR-specific *in vivo* competition defect did not support this model since a strain lacking *rus* in the context of an inability to use fructans was still outcompeted by a strain lacking only fructan use, and this was not due to changes in *rus* activation (**Fig. S4I-J**). Thus, we conclude that the critical role of the Rus ribokinases for *Bt* fitness *in vivo* is contingent upon a more complex set of metabolic interactions that require the generation of a phosphorylated ribose signal(s) that better equips *Bt* to compete in the guts of mice fed the FR diet.

### An enzyme-diversified family of Rus systems exists throughout the *Bacteroidetes*

While the *Bt rus* encodes two enzymes with relatively weak activity and little contribution to the *in vivo* phenotype observed on the FR diet, the architecture of this system suggests that it is equipped to liberate ribose from sources more complex than ribonucleosides. This led us to hypothesize that *rus-*like systems are found in other gut isolates and perhaps more broadly across the *Bacteroidetes* phylum. To test this, we measured the growth ability of 354 different human and animal gut *Bacteroidetes* belonging to 29 species in MM-ribose, revealing that ribose utilization is widely but variably present in indvidual species and strains (**Fig. 6A**, **Fig. S5A**). To determine if sequenced representatives of the species/strains that grow on ribose contain a homolog of the experimentally validated *Bt rus*, we conducted a comparative genomics analysis by searching for homologs of *Bt rus* within these gut isolates. This comparison revealed that all of the sequenced strains with the ability to grow on ribose also possessed a candidate *rus*-like PUL, while none of the strains unable to grow on ribose had a homologous gene cluster.

**Figure 6.**
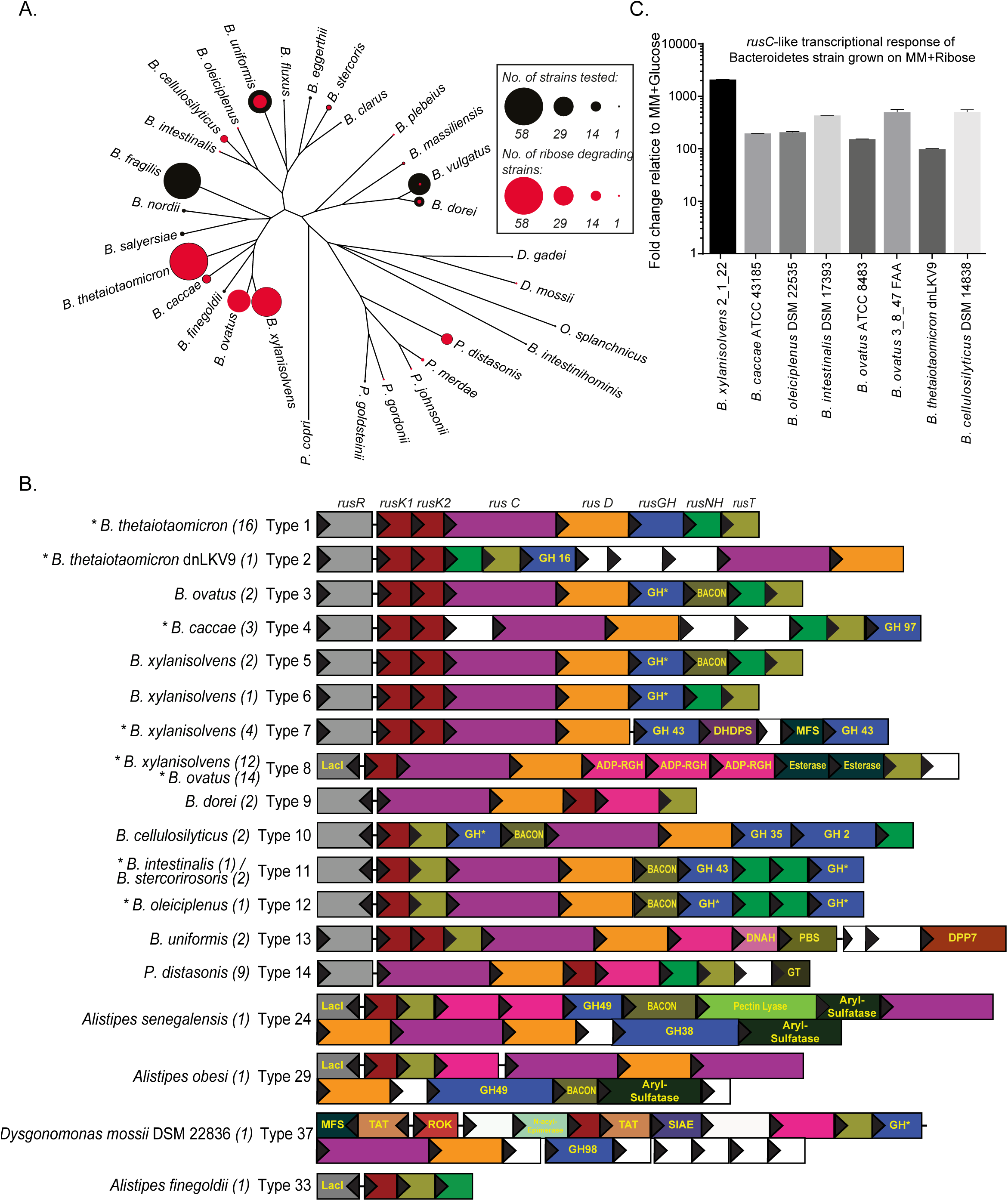
Ribose utilization is present throughout the Bacteroidetes phylum and *rus* homologs appear in many configurations with highly variable enzyme content. (A) Phylogeny of 29 human gut Bacteroidetes species (plus a *P. copri* root) showing the sampling depth of the 354 strains surveyed for growth on ribose and the penetrance of ribose utilization within each species. Outer black circles at tree tips are sized to represent the number of strains sampled within each species, the inner red circles are sized to indicate the number of strains for which ribose (5 mg/ml) supports growth (each strain was tested at least 2 times for growth). (B) Comparative genomics of several variants of homologs of the *rus* locus discovered throughout the Bacteroidetes phylum as described in Methods. The schematic of gene annotations shows the vast enzymatic potential contained in different *rus* loci from the species and strains indicated. Same background fill color indicates the same predicted function(s), no-fill, white backgrounds indicate hypothetical/unknown functions. Adjacent to each locus schematic is the species in which each *rus* homolog is present followed by a parenthetical number denoting the number of sequenced isolates containing the indicated type of *rus* architecture. Each architecture is assigned an arbitrary type number denoting the different gene content and/or organization. All genes are sized relative to actual length within and between genomes. All of the species shown here are human or mammalian gut isolates, a broader representation of *rus* diversity is shown in **Fig. S6** and includes *rus* homologs from environmental and oral Bacteroidetes. Abbreviations: (GH*, Glycoside hydrolase of unknown family/function; BACON, Bacteroidetes-Associated Carbohydrate-binding Often N-terminal domain; DHDPS, dihydrodipicolinate synthase; LacI, predicted lacI-type transcriptional regulator; MFS, Major-facilitator superfamily of transporters; ADP-RGH, ADP-ribosyl glycoside hydrolase; DNAH, DNA helicase; PBS, Polysaccharide Biosynthesis and export of O-antigen and techoic acids; DPP7, Dipeptidyl-Peptidase 7 (serine peptidase); GT, Glycosyl Transferase). For simplicity, gene labels are only shown once with the top-most gene given a label (*i.e*., RusNH in green). However, due to similar colors, genes annotated as BACON, LacI, MFS and aryl sulfatase are labeled throughout). Asterisks next to the organism name indicate that the *rus* homolog type was shown to be upregulated during growth on ribose as the sole carbon source. (C) Fold-change of *rusC*-like transcript from the indicated species during growth on 5 mg/ml of ribose as a sole carbon source compared to growth in MM-glucose. Error bars show the SEM of n=3 biological replicates.

Interestingly, our comparative genomics analysis revealed very similar homologs of some *rus* genes in sequenced gut isolates (*e.g., Prevotella*) that were not tested in our initial survey. When we expanded the search to include these species, plus *Bacteroidetes* isolates found in other body sites and in the environment, we detected *rus*-like systems across the *Bacteroidetes* phylum, with the systems found in the genus *Bacteroides* being most similar to the prototype from *Bt*. Remarkably, we identified a total of 70 different *rus* configurations, ranging from simple two gene units (permease and kinase, which do not formally define a PUL), to *rus* PULs containing as many as 36 genes (**Fig. 6B**, **Fig. S5B**). This analysis revealed that for almost all *rus-*like systems, the following genes are present: *rusC* and *rusD* homologs, an upstream *rusR* homolog or to a lesser extent, different regulator types (LacI is most prevalent after RusR), either one or two ribokinase genes, and a *rusT* homolog. Perhaps most intriguingly, the predicted enzymes found in different *rus-*like systems are exceptionally variable. There are at least 22 different predicted glycoside hydrolase families, along with ADP-ribosylglycohydrolases (ADP-RGH), carbohydrate esterases, nucleoside hydrolases, and other predicted enzymatic activities. This plethora of enzymatic potential encoded in *rus* homologs across the *Bacteroidetes* phylum suggests individual species or strains target different ribose-containing nutrients. To further connect these predicted *rus*-like systems with ribose utilization, we probed the transcriptional response of 8 different systems during growth on MM-ribose, finding that all strains tested exhibited ∼100-1000 fold upregulation relative to a MM-glucose reference (**Fig. 6C**).

## Discussion

Diet impacts the human gut microbiota in many ways and members of the prominent *Bacteroidetes* phylum have developed diverse strategies to liberate sugars from often very complex dietary fiber polysaccharides (Luis et al., 2018; Ndeh et al., 2017). Such abilities equip these bacteria to compete for and utilize dietary and endogenous nutrients to sustain their populations. Diet, microbiome- and host-derived RNA, nucleosides, cofactors and other sources of ribose have been largely unexplored as potential nutrients scavenged by members of the gut microbiota. Our findings not only demonstrate that *Bt* utilizes free- and covalently-linked sources of ribose, but that this metabolic capability contributes to competitive fitness *in vivo* in a diet-dependent fashion—likely through a more complicated metabolic mechanism(s) than just acquisition of ribose. It is also clear from our comparative genomics investigation that the ability to access ribose, probably from diverse sources, extends across the *Bacteroidetes* phylum and is present in many strains from the human gut, oral cavity, and environment.

Based on our results we have developed a working model of ribose utilization built around findings from the *Bt rus* PUL and canonical metabolic pathways (**Fig. 7**). Components encoded directly within this PUL are required for import, recognition and phosphorylation of ribose and presumably more complicated molecules that have yet to be discovered, with RusK1 being important for generating the signal for *rus* induction (**Fig. 5E**). The identity of this inducing molecule is also unknown, but is unlikely to be ribose-5-P, which is presumably made during non-inducing growth on glucose via the PPP to synthesize nucleic acid components and histidine. Data also suggest it may not be ribose-1-P, which is the product of nucleoside phosphorylase and exposure to nucleosides did not activate expression (**Fig. S4G**); although, we cannot rule out that in the absence of a low amount of ribose, nucleosides are not transported to be cleaved by BT4554. This leaves ribose-1,5-diphosphate as a potentially unique candidate. Our model also postulates that the ribokinases are required for catabolism of nucleosides, which are primarily imported and cleaved by the product of the BT4554 phosphorylase. This interconnection may stem from the predicted lack of a dedicated phosphopentomutase in *Bt*, which would be required to convert ribose-1-P generated by BT4554 into ribose-5-P required for entry into the catabolic branch of the pentose phosphate pathway (PPP). As such, at least one additional kinase would be required to create ribose-1,5-PP, a precursor of PRPP that provides an alternative path back to the catabolic branch of PPP, although an orthologous enzyme that catalyzes this transition has also not been identified in *Bt*. In light of a recent study demonstrating that the *Bt* nucleoside phosphorylase (BT4554) has the ability to modify the levels of a nucleoside-containing drug (Zimmermann et al., 2019), our findings hold important implications for how native responses to this overlooked nutrient source can affect the fitness of bacteria and subsequent interactions with the metabolome around them. They also extend the range of enzymatic functions that are encoded in *Bacteroidetes* PULs beyond the known examples of carbohydrate modifying enzymes (Cuskin et al., 2015; Porter and Martens, 2017) and proteases (Nakjang et al., 2012; Renzi et al., 2015) to include those acting on nucleosides.

**Figure 7.**
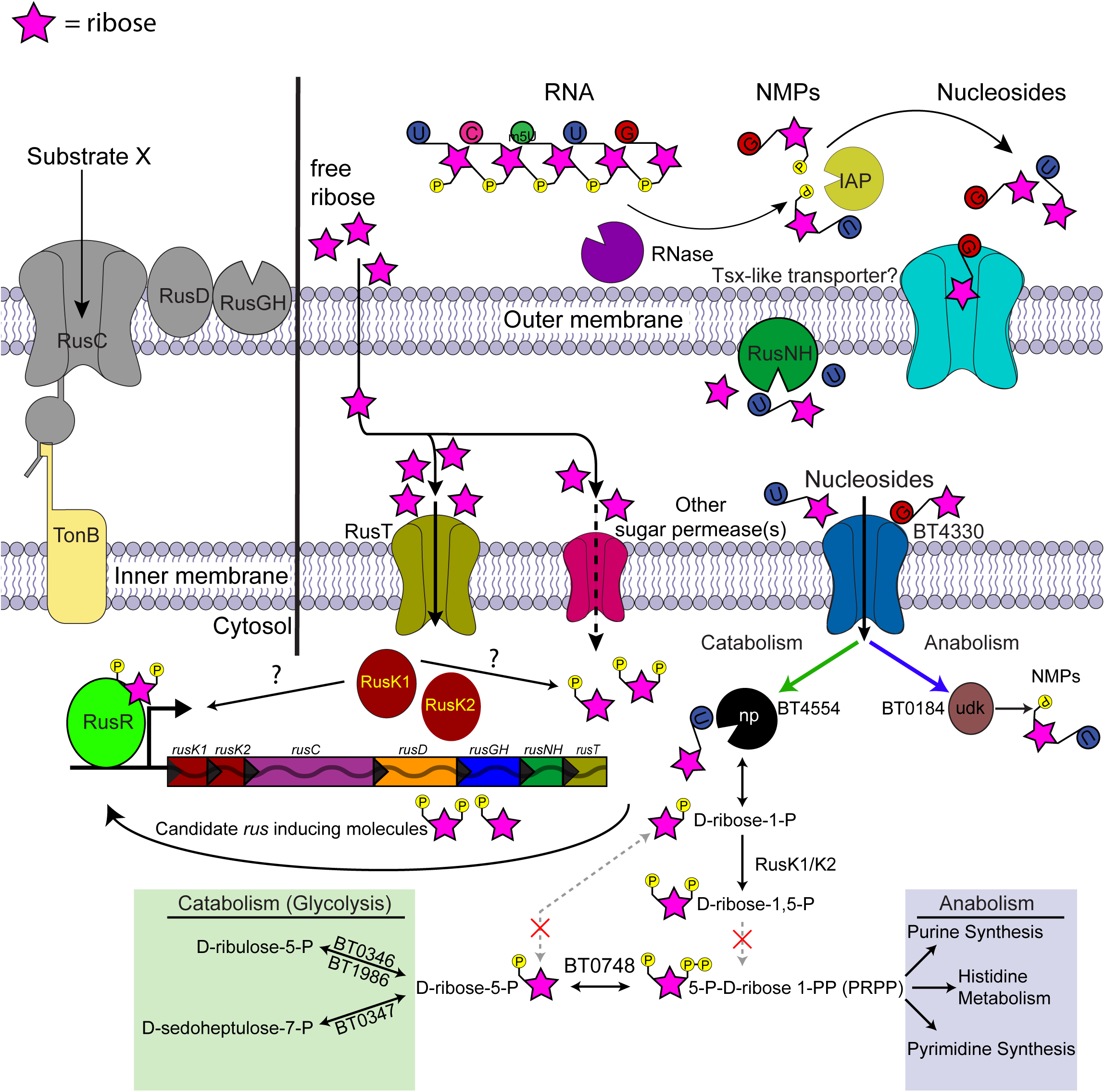
Model of ribose utilization in *Bt*. A schematic of *Bt* metabolism of ribose and related molecules based on the data shown, predicted KEGG metabolism maps of the pentose phosphate pathway, and gene annotations. Ribose is depicted as a pink star, its official symbol nomenclature for glycans symbol, with appended phosphate or bases shown as appropriate (phosphate, yellow; uridine, blue; 5-methyl uridine, green; cytidine, dark pink; guanosine, red). Extracellular ribose should diffuse across the outer membrane and, once in the periplasm, taken up by the high-affinity ribose permease (RusT) or an alternative sugar permease. Transported ribose is then phosphorylated by RusK1 and RusK2 yielding unspecified ribose-phosphates, with the product of at least RusK1 being required for *rus* transcriptional activation. Canonical pathways exist for assimilating ribose-phosphates into either catabolic pathways or synthesis of nucleic acids and histidine, although the precise entry points for RusK1/K2 derived metabolites is unknown, as are the interconversions catalyzed by steps that are not predicted to be present in *Bt* (dashed pathways with red “x” marks). Nucleosides require the additional presence of transport machinery (*e.g*., BT4330) and nucleoside phophorylase activity (*e.g*., BT4554) in order to enter the cell and be catabolized in a *rus*-dependent fashion. Nucleoside transport across the outer membrane in *E. coli* requires a Tsx-like porin, although highly similar candidates were not found in *Bt*.

Similar to only one other PUL that was previously characterized in *Bt*, the prototypic *rus* PUL is activated in response to a monosaccharide (**Fig. 1C**), rather than the more common sensory strategies driven by oligosaccharide cues. The previously described system for fructan utilization is activated by free fructose, which occurs in two common linkages (β2,1 and/or β2,6) in polysaccharides for which various *Bacteroides* strains have acquired substrate-specific enzymes in their respective fructan utilization systems (Sonnenburg et al., 2010). Comparable to the ribose utilization system described here, the fructan system also contains a dedicated permease and a kinase revealing that these two systems are similarly patterned around a core monosaccharide utilization pathway. Taken together, these observations may indicate that one mechanism for PUL evolution and diversification is building enzyme systems, which liberate the same sugar from multiple sources, around a central hub of sugar utilization functions such as import and phosphorylation. While covalently-linked fructose is present in only a few known forms (inulin, levan, graminins) from plant fructans and microbial capsules (van Arkel et al., 2013), ribose is much more widely present but frequently exists in a form linked to a variety of other non-sugar moieties.

Often, the enzymatic content of *Bacteroidetes* PULs provides a window into the fine linkage structure of the nutrient(s) that any given system has evolved to target for degradation. For example, investigation of *Bt* PULs required for degrading α-mannan polysaccharides derived from fungal cell walls revealed enzymes specific for unique modifications in individual fungal species (*Saccharomyces cerevisiae, Schizosaccharomyces pombe* and *Candida albicans*) (Cuskin et al., 2015; Temple et al., 2017). An additional study of xyloglucan utilization PULs across multiple species revealed variants that contain α-fucosidases, presumably equipping them to target fucosylated xyloglucans present in the cell walls of lettuce and other leafy greens (Larsbrink et al., 2014). Ribose is present in many diverse sources with different linkages, including RNA and nucleosides, bacterial capsules, cofactors such as NAD, cellular modification like (poly) ADP-ribose and more exotic molecules such as microcins (Duquesne et al., 2007; Knirel et al., 2002). By coupling ribose monosaccharide sensing, import, and phosphorylation to an adaptable series of enzymes encoding a large repertoire of cleavage functions, we hypothesize that the different ribose utilization systems across the *Bacteroidetes* are tuned to liberate ribose from a wide variety of sources. The breadth of this enzymatic diversity found across the phylum is emphasized by the presence of at least 22 different glycoside hydrolase families, nucleoside cleaving enzymes (nucleoside hydrolases, ADP-ribosylglycohydrolases, NAD-hydrolases), esterases, and more totaling a minimum of 70 different configurations of ribose utilization systems. Interestingly, many prominent *Bacteroides* species isolated from the human gut have ribose utilization systems with similar enzymatic content, suggesting that there may be a narrower range of common nutrient(s) targeted by these organisms in their colonic habitat. However, two of the most complex systems observed also exist in human gut *Bacteroidetes* (*Alistipes senegalensis* and *Dysgonomonas mossi*, **Fig. 6B**), suggesting that they have also adapted to more complex molecules. Finally, the large number of *rus*-encoded glycoside hydrolases suggests that a prominent target of these systems may be ribose-containing polysaccharides used to construct the capsules and exopolysaccharides of other gut bacteria, a dimension of gut microbial ecology and glycobiology that has been poorly explored (Porter and Martens, 2017) despite strains of some common species like *E. coli* using ribose to construct capsules (Hackland et al., 1991).

Since homologs of *rus* are present in numerous members of the human gut microbiota, we hypothesized that at least one source of ribose scavenged *in vivo* would be endogenous RNA from bacteria themselves or turnover of host cells. Consistent with this idea, previous studies demonstrated that *Bt rus* is expressed to high levels *in vivo* during dietary conditions in which fiber is depleted or during *Bt* only chemostat cultures in which biofilms develop and cellular debris including RNA may accumulate (TerAvest, 2013). If an endogenous source of ribose is mostly what is scavenged *in vivo,* competition during low fiber diet feeding should have revealed a bigger defect for the Δ*rus* strain compared to wild-type. However, absence of *rus* did not result in a competitive defect unless ribose was provided in the context of FF diet as a free monosaccharide. Similar supplementation with nucleosides or RNA did not reveal the same effect, suggesting that despite *Bt* being able to utilize nucleosides and enzyme-degraded RNA as carbon sources *in vitro*, dietary sources of these molecules might be absorbed before reaching the distal gut.

Despite lack of identification of a nutrient source(s) that drives the FR-specific diet effect, our study clearly indicates the importance of *rus in vivo*, especially in the FR diet, but also when ribose is present in addition to a low fiber diet. What is more surprising about this result is that monosaccharide analysis of the diets revealed ribose as only a minor constituent of the fiber rich diet in a covalently linked form. One possible mechanism for why the Δ*rus* strain is severely outcompeted in this diet is that utilization pathways for other nutrients may be interconnected with ribose sensing in *Bt* and that small amounts of ribose below the limit of detection, but sufficient to upregulate *rus*, act to enhance utilization of other nutrients *in vivo* or optimize co-utilization of multiple nutrients. This idea is supported by our RNA-seq analysis, specifically the ribose-induced upregulation of genes involved in fructan and arabinose metabolism. Further, several genes responsible for functions encoded in the TCA cycle are upregulated in ribose, suggesting that ribose may remodel the metabolic landscape in such a way to promote faster assimilation of nutrients. Therefore, the ability to sense and respond to ribose via *rus* provides the observed competitive advantage despite only low concentrations of ribose are present from the diet. This phenomenon suggests that nutrients such as monosaccharides should not be overlooked for their intrinsic simplicity and assumption that they do not affect the gut microbiota, as they may still impart changes to individual species’ global metabolism.

A poignant example of how competitive survival in the gut requires evolution of complex nutrient acquisition strategies is exemplified by the *Bt* rhamnogalacturonan-II (RG-II) acquisition system. RG-II consists of 13 different sugars connected through 21 unique linkages and *Bt* contains three co-expressed PULs to scavenge this nutrient, using all of the individual products but one. The results described here highlight how the survival of related bacteria from the human gut and other ecosystems has driven adaptations to sense and scavenge ribose, a ubiquitous sugar that occurs in a number of different molecules, which has apparently led to enormous species and strain level variation in the enzymes present in *rus* loci. This evolution is analogous to a molecular “Swiss-army knife”, in which the core function is utilization of the simple sugar ribose but the various blades and other implements represent the enzymes that equip a given system to sense, import or harvest ribose from one or more sources. This molecular adaptability is particularly important in the context of the nutrient niche hypothesis of gut bacterial survival. While some nutrients may be scarce compared to common and abundant dietary fiber polysaccharides, competition for these lower abundance nutrients may be less intense and organisms capable of accessing them could thereby occupy a stable niche. While a number of gut bacteria, including pathogens, are capable of utilizing free ribose, the *Bacteroides* may have developed a more sophisticated ability to scavenge multiple sources of ribose from covalently linked forms. From this perspective, understanding the struggle to access this “simple” nutrient may reveal additional layers underpinning the interplay between native gut mutualists and invading pathogens.

## Materials and methods

### Bacterial strains, culturing conditions, and molecular genetics

*B. thetaiotaomicron* ATCC 29148 (VPI-5482) and its genetic variants, as well as other *Bacteroides* strains used in this study, were routinely grown in tryptone-yeast extract-glucose (TYG) broth medium (Holdeman, 1977), in minimal medium (MM), plus a defined carbon source (Martens et al., 2008), or on brain heart infusion agar with 10% defibrinated horse blood (Colorado Serum Co.). Unless otherwise noted, carbon sources used in MM were added to a final concentration of 5 mg/ml. Cultures were grown at 37°C in an anaerobic chamber (10% H_2_, 5% CO_2_, and 85% N_2_; Coy Manufacturing, Grass Lake, MI). Genetic deletions and mutations were performed by counter-selectable allelic exchange as previously described (Koropatkin et al., 2008). Primers used in this study are listed in **Table S6** To quantify growth on carbon sources and examine mutant phenotypes, increase in culture absorbance (600 nm) in 200µl cultures in 96-well plates was measured at 10 minute intervals for at least 96 hours on an automated plate reader as previously described (Martens et al., 2011). To achieve consistent and robust growth on nucleosides and other covalently linked sources of ribose, free ribose was added at a final concentration of 0.5 mg/ml to MM containing 5 mg/ml of carbon source. Growth on 5mg/ml of MM containing Type IV Torula yeast RNA (Sigma) was obtained by adding 100 units of calf-intestinal alkaline phosphatase (CIP) (New England Biolabs) and 2mg/ml RNase A (Sigma). Growth parameters and conditions for all substrates are summarized in **Table S1**.

### Genetic manipulation and recombinant protein purification in *E. coli*

To create a nucleoside hydrolase-free expression background, *E. coli* BL21-AI^™^ One Shot^®^ cells (Invitrogen) were manipulated using lambda red recombineering to introduce genetic deletions of the <u>r</u>ibose-inducible <u>h</u>ydrolase genes (*rih*) to avoid contaminating activity in downstream applications of purified proteins (Petersen and Moller, 2001). The *E. coli* gene deletion procedure developed by Datsenko and Wanner (Datsenko and Wanner, 2000) was followed with few modifications. Briefly, BL21-AI cells were transformed with the pKD46 plasmid. Transformed cells were grown overnight in LB + Amp^100^ and sub-cultured, when the culture absorbance (600 nm) reached 0.1, L-arabinose was added to 10 mM final concentration to induce the P_BAD_ promoter of pKD46, cells were allowed to grow to an OD between 0.6-0.8 and made competent for electroporation by cold water washes and stored in 10% glycerol aliquots. For recombineering, 400ng of gel-purified PCR product was added to freshly made cells and incubated for 10 minutes on ice, electroporated in a 2mm gap cuvette at 2500 V, recovered in 1 ml LB at 30°C for 5 hours. All knockouts were made sequentially in this manner via introduction of the following antibiotic cassettes (spectinomycin from K11497 for Δ*rihA*; hygromycin from K11521 for Δ*rihB*; gentamicin from K11590 for Δ*rihC*), and the following concentrations of antibiotic were used for selection: Spec^80^, Hygro^200^, Gent^10^. Following construction of the last deletion, the pKD46 plasmid was heat-cured by passaging twice at 42°C in LB. To better control background expression of the T7 promoter, the T7 lysozyme containing plasmid, pLysS from BL21 (DE3) (Lucigen) was introduced into the strain via Ca^2+^ chemical competence/heat shock. Protein purification was accomplished using the pETite N-His vector (Lucigen). PCR primers were designed to amplify products for BT2807 and BT2808 containing all amino acids downstream of the predicted signal peptide sequences, residues 22-539 for BT2807 and residues 22-338 for BT2808, amplified and transformed into Hi-Control 10G cells according to manufactures protocol (Lucigen, *Expresso*^™^ T7 cloning and expression system). pETite plasmids containing BT2807 or BT2808 were transformed into *E. coli* strains TUNER or BL21-AI Δ*rihABC* + pLysS, respectively. A single colony was grown in 5 mL of LB+Kan^50^ for 16h. This pre-inoculum was added to to 1L of Terrific-Broth with 50ng/ul of Kanamycin and 10 ng/ul of Chloramphenicol (BT2808) or 50ng/ul of Kanamycin (BT2807) and culture was grown with shaking at 37 °C until absorbance 0.4 at 600nm. BT2807 and BT2808 cells were induced with a final concentration of 0.2mM or 1 mM IPTG and 0.2% 20mM L-arabinose, respectively, and temperature was reduced to 16^°^C and outgrown overnight. The recombinant proteins were purified by immobilized metal ion affinity chromatography using cobalt (BT2807) or nickel-affinity (BT2808) columns was accomplished as described previously (Cameron et al., 2014).

### Measurements of transcriptional responses by qPCR

*Bt* and other *Bacteroides* strains were grown to mid-exponential phase 0.6-0.8 (absorbance at 600nm) in MM+ribose, MM+arabinose, MM+xylose, or MM+glucose, two volumes of RNA protect added, followed by centrifugation and storage of cell pellets at −80°C. Total RNA was extracted using the RNeasy mini kit buffers (Qiagen) and purified on RNA-binding spin columns (Epoch), treated with TURBO DNaseI (Ambion) or DNase I (NEB) after elution and purified again using a second RNeasy mini kit isolation column. Reverse transcription was performed using SuperScript III reverse transcriptase and random primers (Invitrogen). The abundance of each target transcript in the resulting cDNA was quantified using either KAPA SYBR® FAST qPCR mix (KAPA Biosystems) or a homemade qPCR mix as described here: each 20 uL reaction contained 1X Thermopol Reaction Buffer (NEB), 125uM dNTPs, 2.5mM MgSO4, 1X SYBR Green I (Lonza), 500nM gene specific or 65nM 16S rRNA primer and 0.5 units Hot Start *Taq* Polymerase (NEB), and 10ng of template cDNA. For the KAPA mix, 400 nM of primers specific for genes in the *rus* locus of *Bt* or the *rusC-*like gene of other *Bacteroides* species or 62.5 nM of 16S rRNA primers and 10ng of template cDNA as described previously (Pudlo et al., 2015). Using the ddCT method, raw values were normalized to 16S rRNA values and then MM+ribose values were referenced to the values obtained in MM+glucose to obtain a fold-change. Measurements of transcriptional response over time in MM+ribose or nucleosides was performed similarly to previously described (Cameron et al., 2014). Briefly, strains were grown in TYG, subcultured 1:50 into MM+glucose, at mid-exponential phase, cells were washed twice in MM-no carbon and resuspended in MM+ribose with time points being taken every 5 min for the first 30 min and every 15 min for a total of 120 min. Measurements of transcriptional responses to varying amounts of ribose were performed similarly as above, but only one time point was taken after 30 min of exposure to varying concentration of MM+ribose ranging from 0.0005 mg/ml to 5mg/ml.

### Gnotobiotic mouse experiments

All experiments involving animals, including euthanasia via carbon dioxide asphyxiation, were approved by the University Committee on Use and Care of Animals at the University of Michigan (NIH Office of Laboratory Animal Welfare number A3114-01) and overseen by a veterinarian. Six to eight-week-old, germfree female Swiss-Webster mice were initially maintained on the standard, fiber-rich lab diet (LabDiet 5010, LabDiet, St. Louis, MO), where appropriate, mice were switched to a fiber-free diet (Envigo-Teklad TD 130343) and maintained for one week prior to colonization with *Bt* strains. After stable colonization had been observed, at day 14 some groups of mice were provided water ab libitum containing one of the following: 1% ribose, 1% Nucleoside mixture (0.25% thymidine, 0.25% uridine, 0.25% 5-methyl uridine, and 0.25% cytidine) or Type VI torula yeast RNA. DNA was extracted from fecal pellets throughout the experiment and strain abundance was quantified as described previously (Desai et al., 2016). Relative abundance of each strain was normalized to the original abundance on day of gavage (day 0). Post-sacrifice, cecal contents were collected, flash frozen and stored at −80°C. RNA was extracted as described previously (Porter and Martens, 2017), briefly, RNA was phenol-chloroform treated and ethanol precipitated, DNA removed by treatment with TURBO^™^ DNaseI (Ambion), followed by purification using RNeasy mini kit (Qiagen) according to manufactures instructions.

### Antibody production, western blotting and immunofluorescent microscopy

Purified recombinant BT2807 and BT2808 proteins were used as antigens to raise rabbit polyclonal antibodies (Cocalico Biologicals, Inc, Stevens PA). Antibody specificity and cellular localization for BT2807 and BT2808 were determined by western blotting of wild-type and relevant mutant strains and by immunofluorescent microscopy of *Bt* VPI-5482 grown in MM+glucose or MM+ribose. Growth conditions are described above, cells for WB were grown to mid-log optical absorbance (600 nm) 0.6-0.7 or 0.4-0.5 for IF. Western blots of *Bt* whole cell lysates were performed using the primary, polyclonal antibodies mentioned above and secondary antibody conjugated to goat anti-Rabbit IgG conjugated alkaline phosphatase (Sigma) and detected with NBT/BCIP (Roche). Surface expression of BT2807 or BT2808 was examined by staining with a BT2807- or BT2808-specific primary antibody in non-permeabilized formaldehyde-fixed *Bt* cells and detected with Alexa-Flour® 488 conjugated goat anti-Rabbit IgG secondary (Molecular Probes), as described previously (Ref). Cells were imaged on an IX-70 inverted microscope (Olympus) with images captured at 100x magnification. A minimum of five fields of view per slide was observed with n=2 biological replicates.

### Comparative genomics of *rus* PULs across Bacteroidetes genomes

A total of 354 different *Bacteroidetes* strains were tested for growth on ribose as a sole carbon source as shown in **Fig. 6A** and summarized in **Table S4** The ability to use ribose is shown in the context of a previously published human gut Bacteroidetes phylogeny that used 14 conserved genes across phylum members (Larsbrink et al., 2014). To search for *rus* locus homologs across the Bacteroidetes phylum, we used the amino acid sequences of the *rusK1, rusK2, rusT,* and *rusR* genes from the *Bt* type strain as deletion of these genes yielded growth defects on ribose. We searched the Integrated Microbial Genomes (IMG) database (current as of May 2018) and performed phylum-level BLAST searches with an E-value cutoff of 1e-50. We chose this stringent cutoff as initial searches using lower values obtained many non-specific hits of genes encoding other kinases and permeases that did not appear to be specific for ribose, including in the *Bt* VPI-5482 genome for which Rusk1 and RusK1 are the only kinases able to promote ribose growth. After we completed our search for *rusK, rusT,* and *rusR* homologs we used the Gene Neighborhood tool in IMG to determine if these hits were located directly next to other genes involved in ribose utilization. The presence of a minimum of two adjacent *rus* gene homologs was required to count the presence of a candidate utilization locus. Following this first round of searching we observed that many of the *rus* loci contained one or more nucleoside cleaving enzymes such as homologs of *Bt rusNH* or ADP-ribosylglycohydrolases (RGH) and upstream putative regulatory genes. To give our search more power and potentially find additional *rus* homologs we performed additional searches with the same E-value threshold for homologs of *Bt rusNH,* or homologs of the ADP-RGH in *B. xylanisolvens XB1A*. When assembling the comparative genomics data, gene names and glycoside hydrolase family assignments are shown as predicted in IMG or by BLAST of the amino acid sequence of individual genes. Further, in refinement, a handful of genes were found below our E-value, but included in the table as it is clear from gene neighborhood views in IMG that it is likely to be part of a *rus* locus. Types of *rus* have been assigned based only on gene content and arrangement as a way to indicate differences, however subtle. In completing our table we have included the bit score as well as the amino acid % identities compared to *Bt rus* genes or *Bx XB1A* ADP-RGH genes. All of the positive gene hits with locus tag information, isolation location, and other relevant strain information is summarized in **Table S5**.

### RNAseq analysis

To determine the global transcriptional response to growth in ribose as the sole carbon source, *Bt* cells were grown overnight in rich TYG media then transferred to fresh MM containing either 5 mg/ml glucose or 5 mg/ml ribose. Cells were then grown until mid-log phase (absorbance between 0.6-0.8) and two volumes of RNA Protect (Qiagen) were added to cells. RNA was isolated as described above and purified whole RNA was then rRNA depleted using the Ribo-Zero Bacterial rRNA Removal Kit (Illumina Inc.) and concentrated with the RNA Clean and Concentrator-5 kit (Zymo Research Corp, Irvine, CA). Samples were multiplexed for sequencing on the Illumina HiSeq platform at the University of Michigan Sequencing Core. Data was analyzed using Arraystar software (DNASTAR, Inc.) using RPKM normalization with default parameters. Gene expression in ribose was compared to gene expression in a glucose reference. Genes with significant up- or down-regulation were determined by the following criteria: genes with and average fold-change ≥5-fold and with at least 2/3 biological replicates with a normalized expression level ≥1% of the overall average RPKM expression level in either glucose or ribose, and a p-value < 0.05 (t test with Benjamini-Hochberg correction) (**Table S3)**.

### Enzyme assays

Recombinant proteins purified in *E. coli*, were used to determine enzyme kinetics for RusGH and RusNH. For RusNH we used a *p-*nitrophenol-ribofuranoside substrate with absorbance readings at 405nm over a 24-hour period as described previously (Desai et al., 2016), with modifications for using purified protein instead of crude extract, using 0.5mM of enzyme in a buffer containing 20mM Hepes and 100mM NaCl, at pH 6.7 at 37°C and continuous absorbance readings. For RusGH, a panel of other 4*-*nitrophenol based substrates in addition to *p-*NP-ribofuranoside were tested at pH 9.0 in 100 mM Tris at 37°C for 16h with 1.5-15 µM of enzyme and using endpoint absorbance measurements. Ion requirements of the RusGH were assayed in *p*-NP-ribofuranoside by addition of divalent cations in the form of CaCl_2_, ZnCl_2_, or MgCl_2_, at 2, 5, or 10 mM concentrations, or in the presence of 10 mM EDTA. Specificity and kinetic parameters for RusNH on natural nucleoside substrates were determined as described previously using a UV-based assay (Parkin et al., 1991). Briefly, a 96-well, UV-compatible microplate (Santa Cruz Biotechnologies) was used with substrate concentrations ranging from 0.025mM-2.5mM, and enzyme concentrations of 0.25-1uM. Assays were immediately read after addition of enzyme by continuous reading of absorbance at 262nm or 280nm with time points taken every 2.5 minutes over 12-24 hours at 37°C. Volume was 250uL in all assays and carried out in buffer containing 20mM Hepes and 100mM NaCl, at pH 6.7, adjusted with acetic acid. As a measure of catalytic efficiency, (*K*_*cat*_/*K*_*M*_) was unable to be determined by classical Michaels-Menton kinetics as Vmax was never reached and therefore Km values were not accurate, so we used a previously established method of estimating this value (Ndeh et al., 2017). Briefly, we used a single substrate concentration to calculate (*k*_cat_/*K*_M_) and checked to be <<*K*_M_ by halving and doubling the substrate concentration and observing a proportionate increase or decrease in rate. Therefore the equation, *V*_0_ = (*k*_cat_/*K*_M_)[S][E] was used to calculate *k*_cat_/*K*_M_ in our case. For, RusGH a panel of other 4*-*nitrophenol based substrates in addition to *p-*NP-ribofuranoside were tested at pH 9.0 in 100 mM Tris at 37°C for 16h with 1.5-15 µM of enzyme with endpoint absorbance measurements. Ion requirements of the RusGH were assayed in *p*-NP-ribofuranoside by addition of divalent cations in the form of CaCl_2_, ZnCl_2_, or MgCl_2_, at 2, 5, or 10 mM concentrations, or in the presence of 1 mM EDTA. The RusGH was tested against a panel of oligosaccharides, nucleosides and nucleotides. Briefly, the reactions were performed with 10 μM of RusGH, 8mg/ml substrate or 5mM monosaccharide in 50 mM TRIS pH 9.0 at 37 °C for 16h. A control reaction was performed in the same conditions without enzyme. The activity was qualitative determined by thin layer chromatography. 6 μl of the reaction was spotted on foil backed silica plate (Silicagel 60, 20 x 20, Merck) and develop in butanol:acetic acid:water 2:1:1 (mobile phase). The products of the reaction were detected by immersing the TLC plate in developer (sulphuric acid/ethanol/water 3:70:20 v/v, orcinol 1 %) for 30 seconds and heating to 100 °C for 2 minutes. A standard of ribose was run in all TLC plates.

### Determination of free and acid hydrolysable monosaccharide content in diets and cecal contents using GC/MS

Prior to analysis, diets were ground to a fine powder using a blender followed by mortar and pestle, while cecal contents were dried by lyophilization. Samples were analyzed for free and linked monosaccharides using the following method described in (Pettolino et al., 2012). In brief, all reactions began with 1-3mg of sample and samples were hydrolyzed in 100ul of 2.5 M TFA for 90 min at 121 °C. Samples were allowed to cool to room temperature (RT) and myo-inositol was added as an internal standard (20ul of 2.5mg/ml) and dried under nitrogen. 150ul of methanol was added, dried and repeated once more. Dried samples were then reduced by dissolving in 50ul of 2M NH_4_OH followed by addition of 50 ul of freshly made 1M NaDB_4_ in 2M NaOH. This mixture was sonicated in a water bath for 1 min, followed by incubation at room temperature for 2.5 hours. 23ul of glacial acetic acid was added and samples dried and evaporated 2x with 250ul of 5% (v/v) acetic acid in methanol, followed by 2x evaporation with 250ul of methanol, drying after each step. Acetylation was done by addition of 250ul acetic anhydrate and sonicated 5 min followed by incubation at 100 °C for 2.5 hours. 2ml of ddH_2_O was added and sample vortexed to dissolve residue, followed by room temperature incubation for 10 min. 1ml of dichloromethane (DCM) was added and vortexed followed by centrifugation at 2000 rpm for 2.5 min. The aqueous phase was discarded and the DCM phase washed 2x with 2 ml of ddH_2_O. DCM phase was dried and residue dissolved in 250 ul acetone. For free monosaccharide analysis the initial hydrolysis step with TFA was not performed. To establish a limit of detection in cecal contents, varying amounts of ribose (0.00002-0.2 mg, in 10-fold increments) were added at the same time as the myo-inositol standard to establish percent recovery throughout the methods used. Acetylated samples were analyzed on a gas chromatography (Agilent Technologies model 7890A) coupled mass spectrometer (Agilent Technologies model 5975C) using a fused silica capillary column (60m x 0.25 mm x 0.2µm SP-2330, Supelco Analytical).

## Acknowledgements

R.W.P.G. was supported by the NIH Molecular Mechanisms of Microbial Pathogenesis Training Program (T32 AI007528) as well as a University of Michigan Rackham Merit Fellowship. We thank the support of the Germ Free Mouse Facility at the University of Michigan for assistance with *in vivo* experiments. This work was supported by National Institutes of Health grants R21-AI-128120, R01-GM-099513, R01-DK-118024 (all to E.C.M).

## Figure Legends

**Figure S1.**
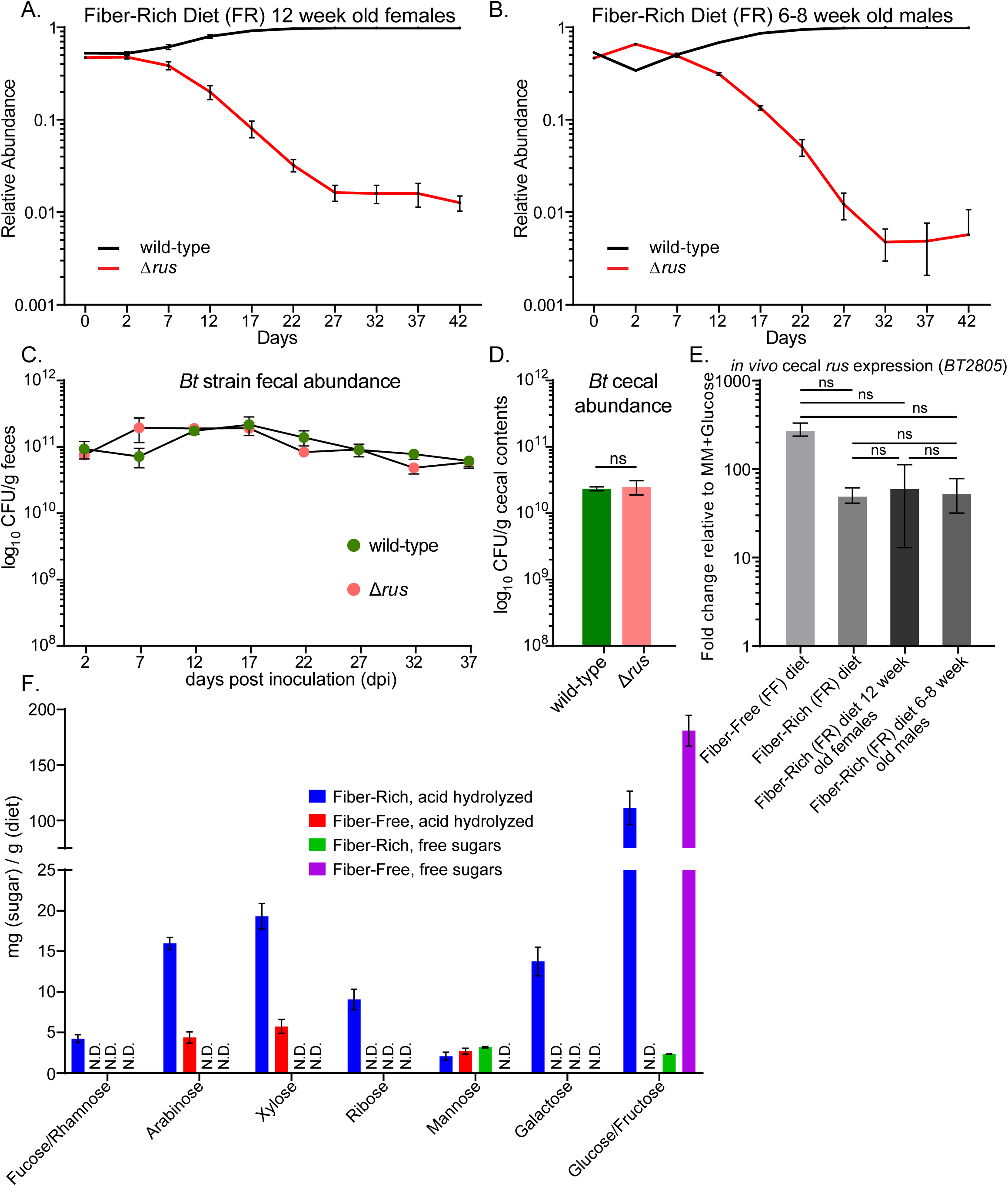
The Δ*rus* and wild-type strain colonization, *rus* expression in all diets, and dietary monosaccharide analysis of FR and FF diets. Data is related to Figure 2. Additional *in vivo* competition or monocolonization experiments performed in Swiss Webster mice fed the fiber rich (FR) diet. Relative abundance of wild type (black) vs. Δ*rus* strain (red) were measured in (A) 12 week old female mice or (B) 6-8 week old male mice. (C) 6-8 week old female mice mono-associated with either wild-type *Bt* (pink circles) or Δ*rus* strain (green circles) and absolute abundance was assayed by dilution plating from fresh fecal samples to obtain CFU/g feces. Data points represent mean plus SEM of n=3 mice. (D) Enumerations of bacterial levels in cecal contents from the mice shown in C. (E) Expression of the *rus* locus in wild-type *Bt* in the cecal contents from experiments in Figure 2A-B (main text) and Figure S2A-B. Data points represent mean plus SEM of n=4 mice. Student’s t-test was used to compare all other samples to *rus* expression in 6-8 week old females on FR diet revealing no significant (ns) differences. (F) GC/MS analysis of free and linked (acid-hydrolyzable) monosaccharides in the FR and FF diets. Data is presented as mg of sugar per gram of diet. Bars show the average plus SEM of n=3 technical replicates. Note that the FF diet contains 44% w/w glucose, which is the source of the strong free glucose signal observed with that diet.

**Figure S2.**
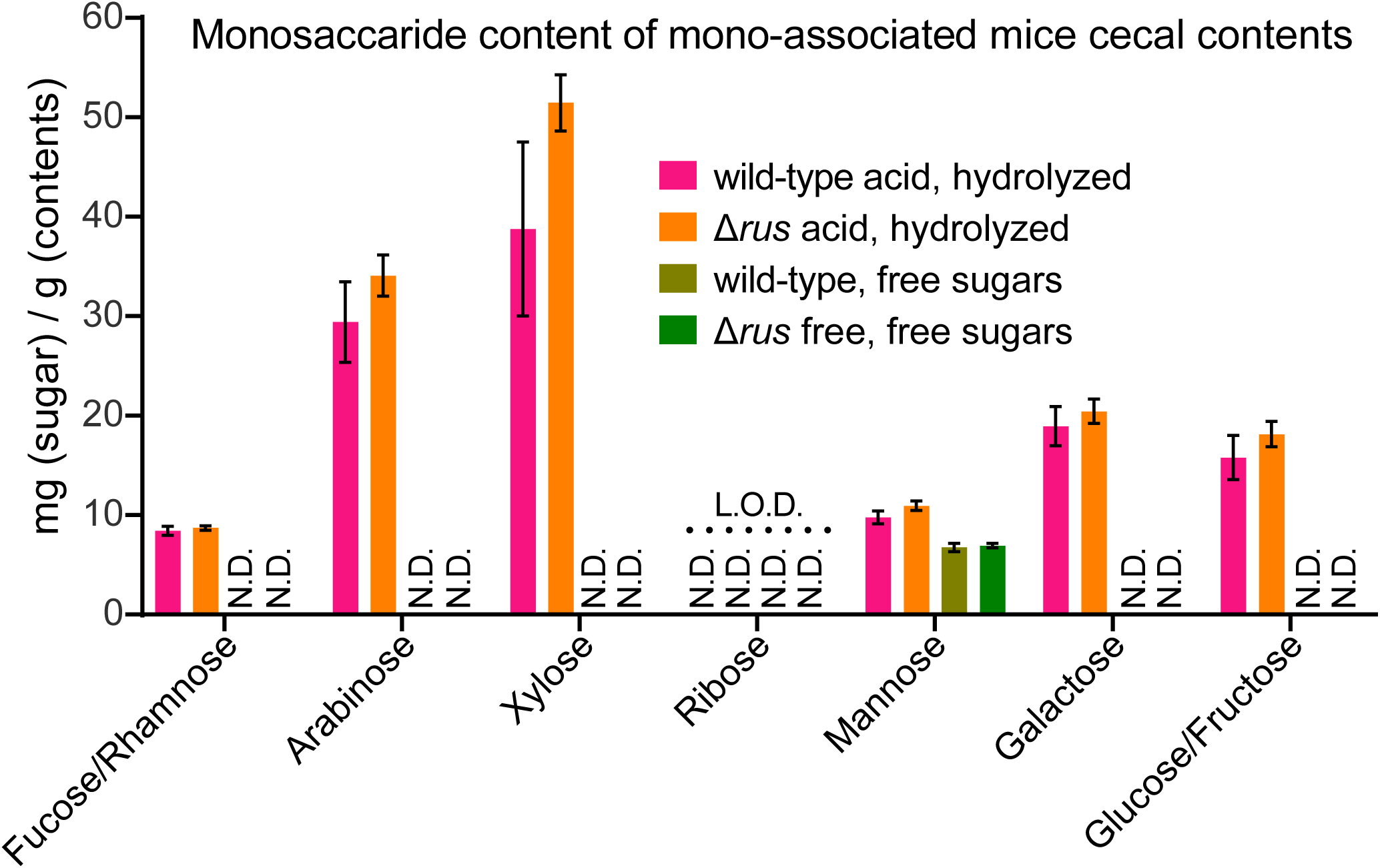
Monosaccharide content of mono-associated mice from cecal contents. Data is related to Figure 2. GC/MS analysis of free and linked (acid-hydrolyzable) monosaccharides from the cecal contents of wild type or Δ*rus* mono-associated 6-8 week old, female Swiss Webster mice maintained on a fiber rich diet for 42 days. Data is presented as mg of sugar per gram of cecal contents. The pink and olive green bars represent wild type linked and free respectively, while the orange and green represent linked or free for the Δ*rus* colonized mice. Bars show the average plus SEM of n=3 technical replicates. N.D. indicates that the sugar was not detectable above our limit of detection (L.O.D.) which is shown as a dotted line above the ribose bars.

**Figure S3.**
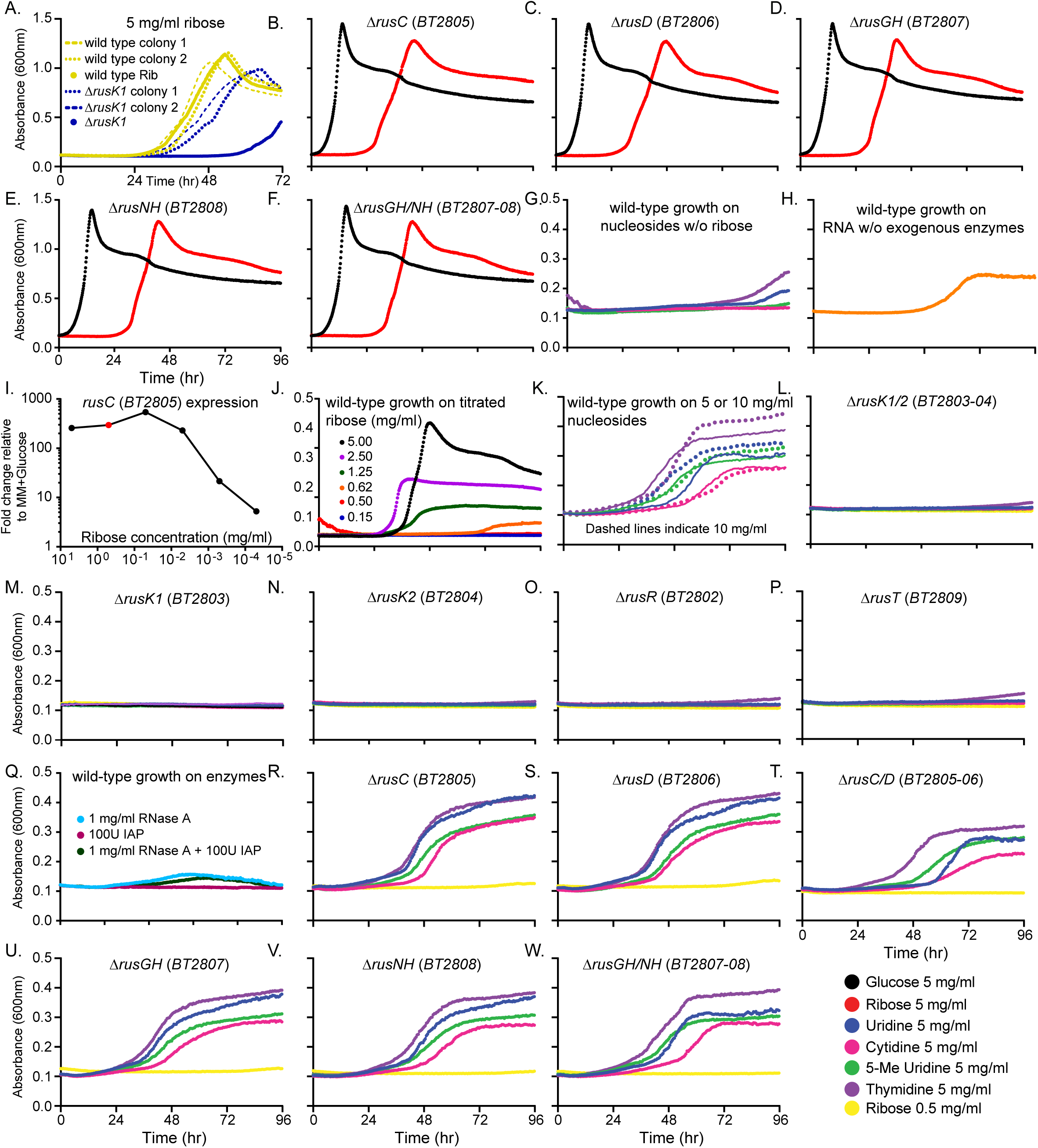
*Rus* deletion strains on ribose containing compounds. Data is related to Figure 4. (A) Wild-type (maize, solid line) or Δ*rusK1* (blue, solid line) strains grown in MM-ribose compared to wild-type and Δ*rusK1* strains that had been previously grown on MM-ribose, isolated on solid medium (BHI-blood), two separate colonies picked into rich media (TYG), and then grown in MM-ribose again shown as maize dashed lines (wild type) or blue dashed line (Δ*rusK1*) to check if the delayed growth phenotype associated with the Δ*rusK1* strain was the product of a genetic suppressor mutation or similar epigenetic/reprogramming. (B-F) Growth of *rus* deletion strains exhibiting similar or identical growth as wild-type on MM-ribose (red) with no obvious growth defects. Growth on MM-glucose (black) is shown for comparison. (G) Wild-type *Bt* grown in MM containing 5 mg/ml of one of the following nucleosides (uridine, blue; cytidine, pink; 5-methyl uridine, green; or thymidine, purple) without any ribose added. (H) Wild-type *Bt* grown on MM containing 5 mg/ml of yeast RNA without any exogenous enzymes. (I) *rusC* transcriptional response when wild-type *Bt* was exposed to titrated amounts of ribose (mg/ml) where each data point represents a different 10-fold dilution of ribose. The red dot represents 0.5 mg/ml ribose which induces *rus* activation to comparable levels as 5 mg/ml ribose the data point directly to the left of the red point compared to growth in MM-glucose. (J) Wild-type growth on different concentrations of MM containing ribose at the following concentrations (mg/ml): 5, black; 2.5, purple; 1.25, green; 0.625, orange; 0.5, red; or 0.15, blue. Growth was not detectable at levels ≤0.5 mg/ml. (K) Wild-type growth on MM containing nucleosides at concentrations of 5 mg/ml (solid lines) or 10 mg/ml (dashed lines) in the presence of 0.5 mg/ml ribose. Individual nucleoside growths are colored same as (G). (L-P) *rus* deletion strains exhibiting a complete lack of growth phenotype on all nucleosides tested are shown as labeled. (Q) Wild-type *Bt* does not grow detectably in MM containing the enzymes used for RNA degradation. (R-W) Individual mutants (as labeled) that do not show a growth defect on nucleosides in the presence of 0.5mg/ml ribose.

**Figure S4.**
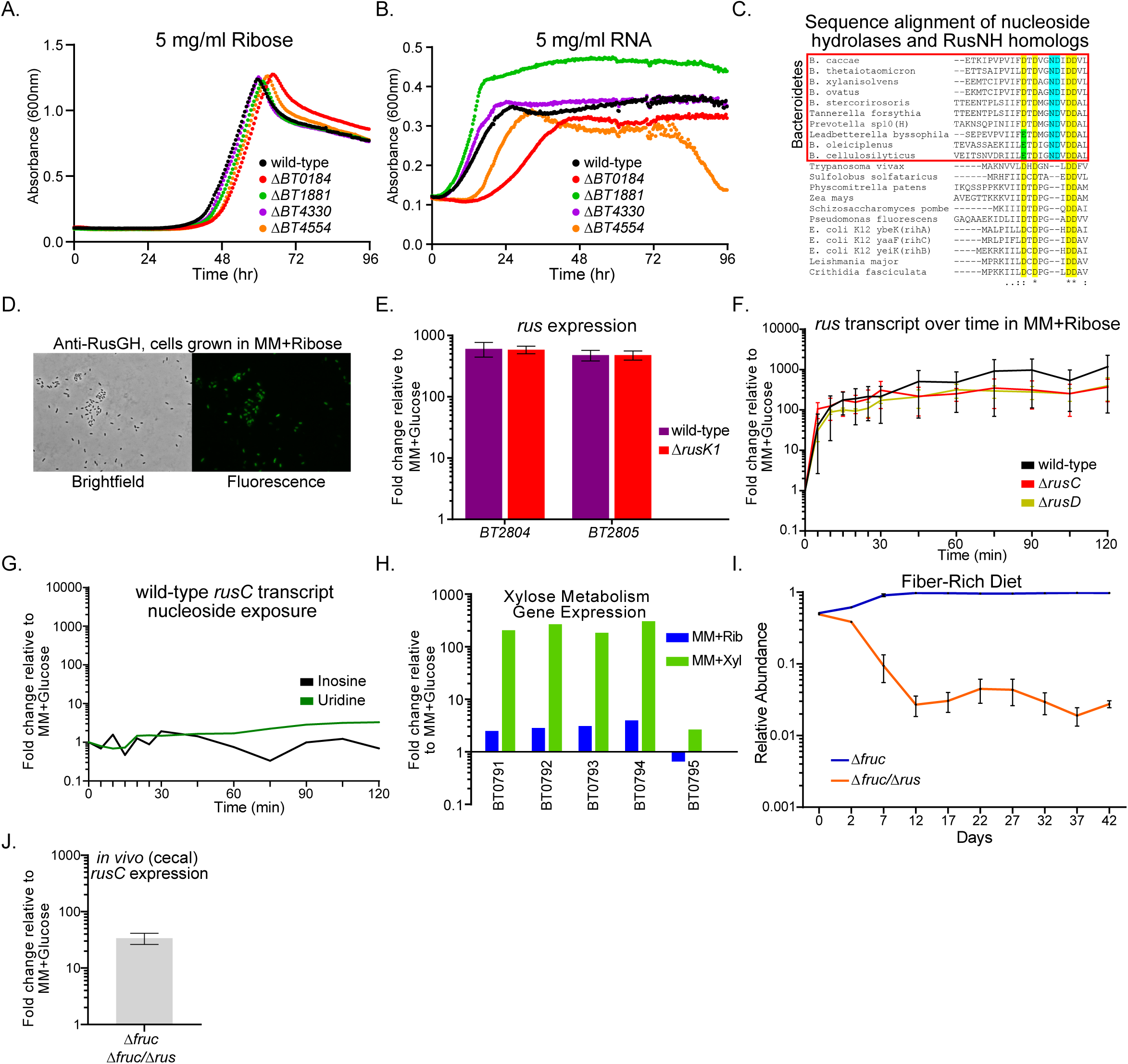
Further characterization of nucleoside scavenging mutants and *rus* transcript activation dynamics, and relevance of global gene response to growth on ribose. Data is related to Figure 5. (A,B) Growth of NSS mutants or wild-type *Bt* on ribose (A) or RNA (with added enzymes). Strains are color coded (wild-type, black; Δ*BT0184*, red; Δ*BT1881*, green; Δ*BT4330*, purple; or Δ*BT4554*, orange). (C) Multiple sequence alignment of BT2808 (RusNH) and other RusNH-like proteins from Bacteroidetes (red boxed region) compared to previously validated nucleoside hydrolases isolated from bacteria (*E. coli,* RihA,B,C and *P. fluorescens*), archaea (*S. solfataricus*), parasitic eukaryotes (*T. vivax, L. major,* and *C. fasciculata*), moss (*P. patens*), maize (Zea mays), and yeast (*S. pombe*), indicating that the predicted nucleoside hydrolase of BT2808 shares the universally conserved N-terminal DXDXXXDD motif responsible for Ca^2+^ coordination (2^nd^ and 4^th^ yellow-highlighted aspartic acid residues) and ribose binding (3^rd^ yellow-highlighted aspartic acid), as well as the nearly conserved canonical 1^st^ aspartic acid residue denoting the motif (yellow or green highlighted position). Specific to the Bacteroidetes nucleoside hydrolases, there are two additional conserved residues, an asparagine and an additional aspartic acid within the motif (highlighted in teal) not found IUNH family nucleoside hydrolases outside of the Bacteroidetes. (D) Immunofluorescent microscopy of intact *Bt* whole cells grown in MM-ribose media staining with anti-BT2807 (RusGH) antibody, indicating that the protein is localized to the outer surface as the secondary antibody has clearly labeled nearly all of the cells seen in the brightfield image (left), a green color in the fluorescent image at (right). (E) Expression of *BT2804* and *BT2805* transcript in wild-type *Bt* (purple) or Δ*rusK1* (red) strains after active growth had initiated in MM-ribose and cells were sampled in mid-log phase and compared to growth in MM-glucose. Bars represent mean, plus SD of n=3 replicates. (F) *rus* transcript during a time course experiment in which cells were shifted from growth on MM-glucose to MM-ribose and transcript measured over time with points every 5 minutes for the first 30 minutes and every 15 minutes after out to the conclusion at 120 minutes post-ribose exposure. For the wild-type (black) and Δ*rusD* (dark yellow) strains, the *rusC* gene was probed, while for the Δ*rusC* strain (red), the *rusD* gene was probed (similar kinetics were seen in the wild-type strain when the *rusD* gene was used to assay *rus* activation, data not shown). Error bars represent the SEM of n=3 replicates performed on separate days. (G) Similar to the response experiment in Figure 5E but using nucleosides (inosine, black line or uridine, green line) and probing *rusC* expression to address if either a catabolized (uridine) or non-catabolized (inosine) ribose containing compound could stimulate *rus* activation, with no response detected compared to growth in MM-glucose. (H) As in Figure 5F, we probed expression of the genes required for xylose metabolism (*BT0791-0795*) when wild-type *Bt* was grown on MM-xylose (green bars) or MM-ribose (blue bars) and compared to *Bt* grown in MM-glucose. (I) *in vivo* competition of a strain lacking the entire fructan PUL, *BT1754-1765* (Δ*fruc*, blue line) against a strain lacking both the fructan PUL and the *rus* PUL (Δ*fruc*/Δ*rus*, orange line) in 6-8 week old Swiss-Webster female mice maintained on the FR diet. The relative fecal abundance is shown on a log scale as assayed by qPCR over the course of the experiment, error bars show the SEM of n=4 mice. (J) *in vivo rus* expression from cecal contents of the mice in I. measuring expression of the *rusC* gene, error bar shows the SEM of n=4 mice.

**Figure S5.**
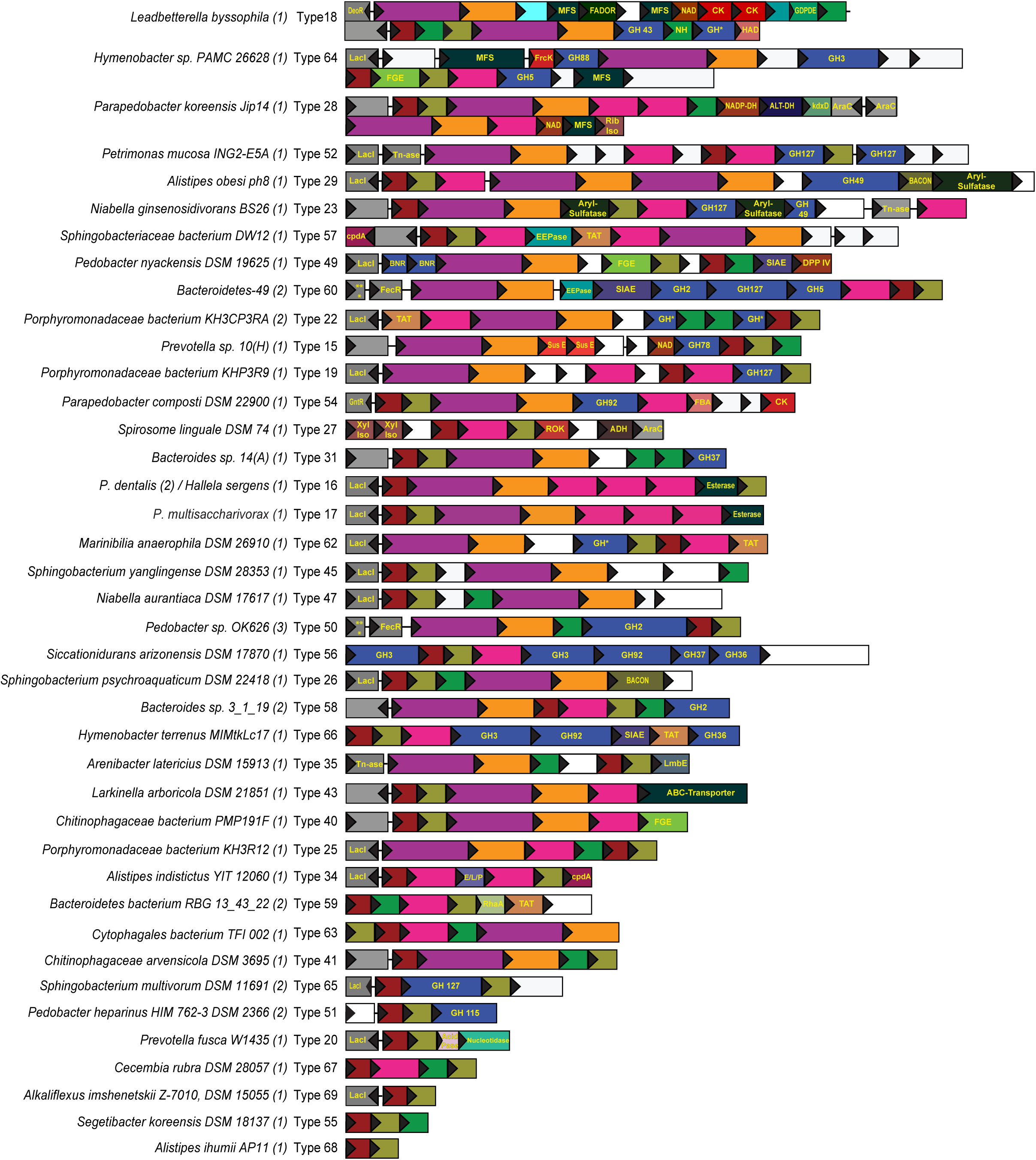
An expanded repertoire of *rus* architectures across the Bacteroidetes phylum. Related to Figure 6. Comparative genomics analysis of a broader survey of members of the Bacteroidetes phylum, revealing many different types of the *rus* locus. This figure displays almost all of the additional locus types found in both the human gut isolates along with those found in aquatic, soil, and human oral environments. Not shown are subtypes, where the same genes are present, but arranged differently, as well as types 21, 30, 33, 36, 39, 44, 48, 46, and 61, for which only one example was identified and gene arrangements had less complexity than the majority shown here (all loci are listed in **Table S5)**. As in Figure 6, the gene size is scaled within and between genomes and the background color is kept constant for genes predicted to encode the same functions. Gene abbreviations are as follows in order of appearance: (DeoR, DeoR-like family of transcriptional regulator; MFS, Major Facilitator Superfamily of transporters; FADOR, Flavin (FAD) Oxidoreductase; NAD, NAD Binding Protein; CK, Carbohydrate Kinase, unknown family; GDPDE, Glycerophosphoryl Diester Phosphodiesterase; NH, Nucleoside Hydrolase; GH, Glycoside hydrolase; HAD, Haloacid Dehydrogenase; LacI, LacI-type transcriptional regulator; FrcK, fructokinase; FGE, Formylglycine-Generating Enzyme, required for sulfatase activity; NADP-DH, NADP-Dependent aldehyde Dehydrogenase; ALT-DH, Altronate Dehydrogenase; kdxD, 2-dehydro-3-deoxy-D-arabinonate dehydratase; AraC, AraC-like transcriptional regulator; Rib Iso, Ribose-5-Phosphate Isomerase; Tn-ase, transposase; BACON, Bacteroidetes-Associated Carbohydrate-binding Often N-terminal domain; cpdA, 3’.5’-cyclic AMP phosphodiesterase; EEPase, Endo-Exo Nucleoside-Phosphatase; TAT, Twin-Arginine Translocase; BNR, BNR repeat-like domain; SIAE, Sialate O-acetylesterase; DPP IV, Dipeptidyl-peptidase IV; ***, RNA polymerase sigma factor ECF subfamily; FecR, FecR-like transcriptional regulator; SusE, Bacteroides SusE-like outer membrane binding protein; GntR, GntR-like transcriptional regulator; FBA, Fructose Bisphosphate Aldolase; Xyl Iso, Xylose Isomerase; ROK, Repressor/ORF/Kinase domain containing protein; ADH, Alcohol Dehydrogenase; LmbE, N-acetylglucosaminyl deacetylase LmbE-like family; E/L/P, Esterase/Lipase/Peptidase-like domain containing protein; RhaA, Regulator of RNaseE activity; Acid Pase, Acid Phosphotase-like protein).

